# Blockwise Site Frequency Spectra for Inferring Complex Population Histories and Recombination

**DOI:** 10.1101/077958

**Authors:** Champak R. Beeravolu, Michael J. Hickerson, Laurent A.F. Frantz, Konrad Lohse

**Affiliations:** Biology Department, The City College of New York, New York, NY 10031, USA; The Graduate Center, The City University of New York, New York, NY 10016, USA; Division of Invertebrate Zoology, American Museum of Natural History, New York, NY 10024, USA; Paleogenomics and Bio-Archaeology Research Network, Research Laboratory for Archeology and History of Art, University of Oxford, Oxford OX1 3QY, UK; Institute of Evolutionary Biology, University of Edinburgh, King’s Buildings, Edinburgh EH9 3FL, UK

**Author notes:** Present address: Institut für Evolutionsbiologie und Umweltwissenschaften, Universität Zürich, 8057 Zürich, Switzerland (Champak R. Beeravolu).

## Abstract

We introduce ABLE (Approximate Blockwise Likelihood Estimation), a novel composite likelihood framework based on a recently introduced summary of sequence variation: the blockwise site frequency spectrum (bSFS). This simulation-based framework uses the the frequencies of bSFS configurations to jointly model demographic history and recombination and is explicitly designed to make inference using multiple whole genomes or genome-wide multi-locus data (*e.g.* RADSeq) catering to the needs of researchers studying model or non-model organisms respectively. The flexible nature of our method further allows for arbitrarily complex population histories using unphased and unpolarized whole genome sequences. *In silico* experiments demonstrate accurate parameter estimates across a range of divergence models with increasing complexity, and as a proof of principle, we infer the demographic history of the two species of orangutan from multiple genome sequences (over 160 Mbp in length) from each species. Our results indicate that the two orangutan species split approximately 650-950 thousand years ago but experienced a pulse of secondary contact much more recently, most likely during a period of low sea-level South East Asia (∼300,000 years ago). Unlike previous analyses we can reject a history of continuous gene flow and co-estimate genome-wide recombination. ABLE is available for download at https://github.com/champost/ABLE.

## Introduction

Demographic history has a major role in shaping genetic variation. However, using this information in an efficient way to infer even very simple models of population history remains challenging: a complete description of the history of genomic samples includes both the ancestral process of coalescence and recombination, as captured by the ancestral recombination graph (ARG). While the ARG is straightforward to simulate, in practice, the number of recombination and coalescent events in any stretch of genome generally exceeds the information (i.e. number of mutations) available to reconstruct them. Thus, it is currently not feasible to perform demographic inference by integrating over all realizations of the ARG that are compatible with a genomic dataset [1].

Current methods dealing with genomic data tackle this problem by making simplifying assumptions about recombination [2]. Methods based on single nucleotide polymorphisms (SNPs) ignore linkage information altogether and make use of the site frequency spectrum (SFS) [3,4], which is a function only of the expected length of genealogical branches [5,6]. While the computing (or approximating) likelihoods based on the SFS is very fast, much of the information about past demography is sacrificed and recent studies have shown that different demographic histories can give rise to a similar SFS [7].

Other methods seek to use linkage information by approximating recombination, i.e. the sequential transitions between local genealogies along the genome, as a Markov process [8,9]. Methods based on the Sequential Markov Coalescent (SMC, [10]) are computationally intensive, limited to relatively simple models [11] or small samples [8,12,13] and require good genome assemblies which are presently available only for a handful of species.

*Multi-locus* methods exploit information contained in short-range linkage by assuming that recombination has is negligible effects within short blocks of sequence [14–18]. However, this approach potentially biases demographic inference and still looses information contained in longer range linkage disequilibrium (LD), which is expected to result from historical admixture or drastic changes in population size. While recombination within blocks has been included in multi-locus inference, this currently does not scale to whole genome data [19]. Interestingly, the few methods capable of jointly inferring recombination (using the SMC) and demography using whole genomes [12,13] can only analyze a couple of samples or are restricted to specific population histories [32,33].

To overcome these limitations, we introduce a composite likelihood (CL) framework which is highly flexible both in terms of the demographic histories and data that can be accommodated. We can infer arbitrarily complex demographic histories along with the average recombination rate using multiple whole genomes or genome-wide multi-locus data (*e.g.* RADSeq) catering to the needs of researchers studying model or non-model organisms respectively. Our method builds upon an existing analytic approach [16,18] that partitions the genome into blocks of equal (and arbitrary) size and summarizes the genome-wide pattern of linked polymorphism as a frequency distribution of blockwise site frequency spectra. We refer to this straightforward extension of the SFS, as the distribution of blockwise SFS configurations, or simply the bSFS. The bSFS is a richer summary of sequence variation than the SFS which retains information about the variation in genealogies contained in linkage within blocks. We use Monte Carlo simulations from the coalescent with recombination to approximate the bSFS. This overcomes the limitations of exact likelihood calculations [18,20] based on the bSFS by accommodating larger samples of genomes and including recombination within blocks as a free parameter. Our approach is implemented in the software Approximate Blockwise Likelihood Estimation (ABLE) which is freely available (https://github.com/champost/ABLE).

The paper is structured as follows: we first describe how the bSFS can be approximated for samples from single and multiple populations both with and without recombination. The accuracy of our approximation is assessed by comparing it to analytic results for small samples in the absence of intra-block recombination under three different demographic models. We then illustrate the performance of ABLE on real data by analysing whole genomes of the two species of orangutan (*Pongo pygmaeus* and *P. abelii*) which inhabit the islands of Borneo and Sumatra respectively [21,22]. These sister taxa represent an excellent test case as their demographic history has been the subject of several previous analyses [12,13,19,21–23] and the geological knowledge of the Sunda shelf is extensive [24]. The best supported history we infer is a previously unexplored scenario of a population divergence (about a million years ago) followed by a discrete pulse of bidirectional admixture which coincides with a cyclical sea-level changes in South East Asia [24]. We also obtain plausible estimates for the per generation genome-wide recombination rate. Finally, we make use of extensive simulations to asses the inferential power of our approach. We explore the ability of ABLE to distinguish between various two population models and investigate the effects of sample and block size on parameter estimates. We also compare the performance of a *small-sample* inference with ABLE to that based on the SFS (∂a∂i [3]) using larger samples.

## Results

### The blockwise SFS (bSFS)

Consider a random sample of sequence blocks of fixed length. In practice, such sequence blocks (coloured segments in Fig. 1) may be obtained by partitioning an available reference genome [20,25] or from reduced representation sequencing strategies, such as restriction site associated DNA (RADSeq, [26]).

**Fig 1.**
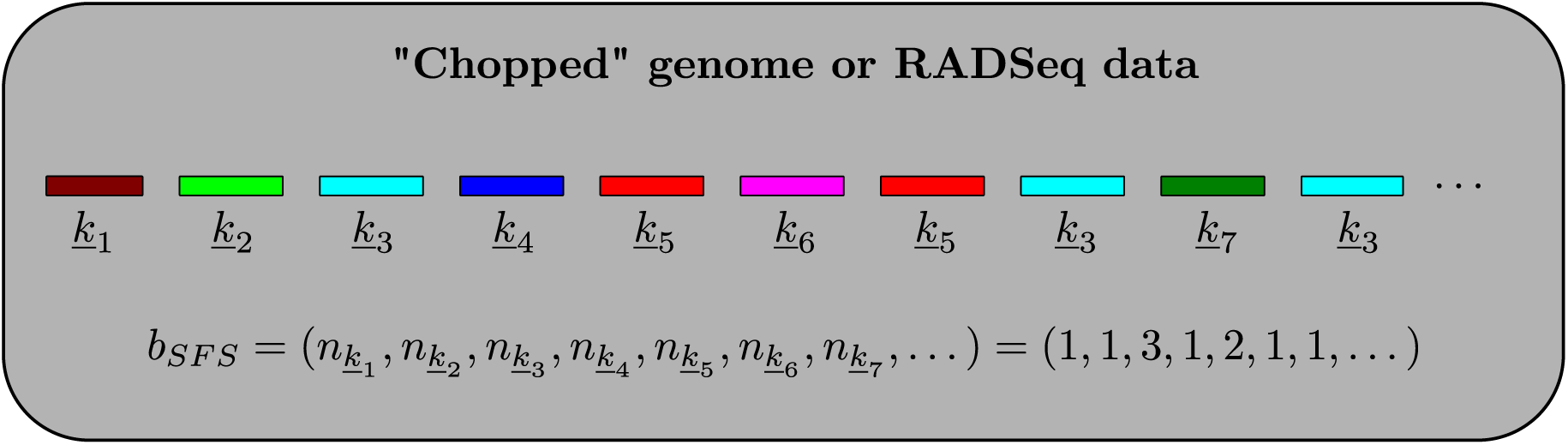
The blockwise SFS (bSFS). The bSFS is computed by partitioning sequences into short blocks, identifying mutation configurations (see Fig. S1) and noting their respective counts.

Given a sample of *b* genomes, the polymorphic sites in each sequence block can be summarized by a vector <u>*k*</u> of length *b -* 1 (Fig. S1). For a single panmictic population, <u>*k*</u> is the SFS of the block and summarizes polymorphic sites within it as counts of singletons, doubletons *etc.* Following [20], the bSFS is essentially a frequency spectrum of site frequency spectra types across blocks (i.e. a histogram of histograms) and can be thought of as a straightforward extension of the SFS that accounts for linkage over a fixed length of sequence block (Fig. 1).

The bSFS readily extends to samples from multiple populations where the entries of <u>*k*</u> are counts of mutation types defined by the joint SFS [6]. One advantage of the bSFS is that we only require *unphased* data as mutations are not distinguished based on unique branches but branch classes (singletons, doubletons, *etc.*, see Fig. S1). In the absence of outgroup information and/or to avoid biases due to errors when polarizing with distant outgroups, the bSFS may be *folded*. The analytical treatment of Lohse *et. al.* [18] (see also [20]) assumes non-recombining blocks and uses a recursion for the generation function of genealogies to derive the probability of bSFS configurations for small samples and simple demographic histories involving one or two populations. This allows for a direct comparison with the approximate composite likelihood developed here.

### Approximating the bSFS

The bSFS can be approximated for any given population history while accommodating for intra-block recombination (see Methods). In summary, we use coalescent simulations corresponding to the size of sequence blocks and compute analytically the probability of observing all bSFS configurations in the data conditional on a particular simulated ARG. Thus, each simulation replicate contributes to the approximate likelihood of all configurations compatible with it. We used a two step optimization procedure to hone in on the Maximum Composite Likelihood Estimate (MCLE) for a given demographic model (see Methods).

### Comparison with analytic results

To study how the number of sampled ARGs summarized by the bSFS affects the convergence of the approximate CL to the analytical expectations, we considered small samples under three simple demographic models: a single population (*b* = 4, no outgroup) which doubled in effective size (*N*_*e*_) at time *T* = 0.2 (Fig. 2a), a history of isolation between two populations A and B (at time *T* = 1.2) followed by continuous unidirectional migration (IM) at a rate *M* = 4*N*_*e*_*m* = 0.5 migrants per generation from A to B (*b* = 2 per population, no outgroup, Fig. 2b) and a history of isolation between three populations (*b* = 1 per population with outgroup) with a recent instantaneous and unidirectional admixture (IUA) that transfers a fraction *f* of lineages from population A to B (Fig. 2c). Parameters under the latter model were chosen to correspond roughly to the divergence and admixture history of humans and Neandertals: *f* = 0.6, *T*_2_ = 0.6, *T*_1_ = 0.15, *T*_*gf*_ = 0.125 [25]. All times are measured in 2*N*_*e*_ generations. For the sake of simplicity, the models in Figs. 2a,b assume identical *N*_*e*_ for all current and ancestral populations (see also [27,28]). The analytic solution for the bSFS under these models were previously obtained using an automation for the generating function implemented in *Mathematica* [18,20,25].

**Fig 2.**
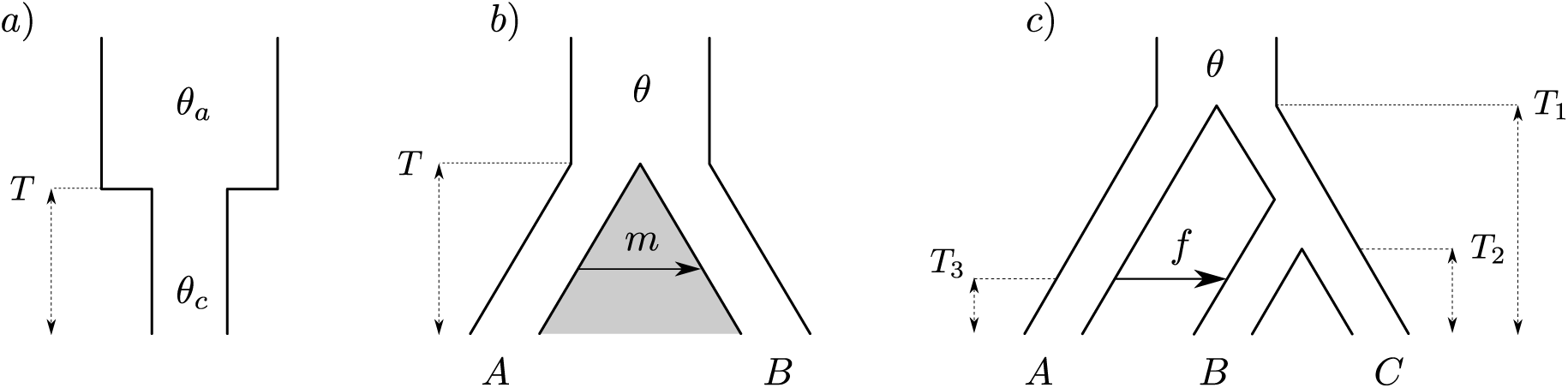
Three demographic models for which ABLE was compared against analytic expectations for the bSFS. (*a*) a single population with a sudden reduction in *N*_*e*_, (*b*) IM:isolation between populations A and B followed by continuous unidirectional migration (from A to B) at rate *M* migrants per generation and (*c*) IUA: isolation between three populations A, B and C followed be unidirectional admixture of a fraction *f* from A to B. Analytical expectations for these models can be found in [18,20,25].

The Monte Carlo approximation to the distribution of bSFS configurations matches the analytic prediction extremely well (Fig. 3) even when only small samples of genealogies are used, e.g. 1000 simulated replicates. This is perhaps surprising, given that this sample size is on the same order as the number of unique bSFS configurations. For example, for a sample of *b* = 2 from the two populations IM model (Fig. 2b) and counting up to *k*_*max*_ = 4 mutations per SFS-type and block, there are 396 unique bSFS configurations. Interestingly, the probability of bSFS configurations involving fixed differences (Fig. 3; green middle row) can be approximated accurately with fewer sampled genealogies than the probability of rare configurations that include shared polymorphism (Fig. 3; blue middle row). The latter are due to either incomplete lineage sorting or admixture. Likewise, for the IUA model, the probability of bSFS configurations involving mutations shared by A and B was harder to approximate than that of (*B,* (*A, C*)) configurations (blue vs. magenta in Fig. 3, bottom row).

**Fig 3.**
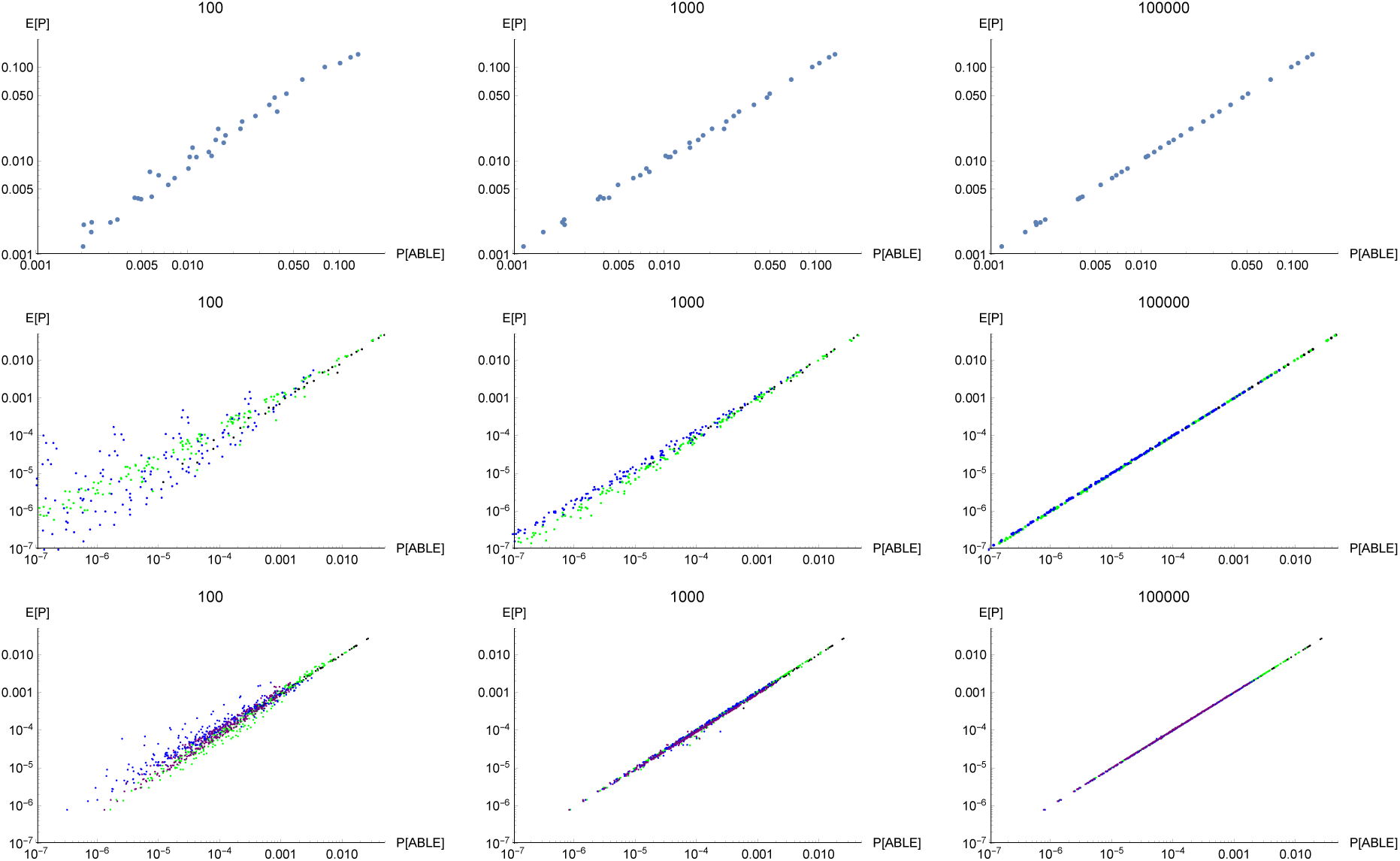
ABLE converges to the expected analytic probabilities with increasing numbers of simulated genealogies (100, 1000 and 10000). Results are shown for models specified in Figs. 2a,b c (top, middle and bottom row) assuming blocks of length θ = 0.6, 1 and 2.4 respectively. For the IM model (Fig. 2b) (middle row) bSFS configurations with shared polymorphisms are shown in blue, those involving fixed differences in green, those with neither in black. For the IUA model (Fig. 2c, here in the bottom row) blocks with topology (*A,* (*B, C*)), (*C,* (*A, B*)) and (*B,* (*A, C*)) are shown in green, blue and magenta respectively. Topologically uninformative blocks are in black.

## Orangutan analyses

To demonstrate the performance of our new bSFS approach using the ABLE framework on real data, we re-analysed whole genome data [21,22] for the two species of orangutan (*Pongo pygmaeus* and *P. abelii*) which inhabit Borneo and Sumatra respectively. These sister taxa are an excellent test case given that their demographic history has been the subject of several previous analyses [12,13,19,21–23]. We selected a subsample consisting of two diploid genomes per species (*i.e. b* = 4 per island) and partitioned the entire autosome into blocks of 2kb (on average 8.22 SNPs/block). After filtering, a total length of 163 Mb of sequence was retained in the final dataset (see Data processing for details), which consisted of 36,544 unique bSFS configurations. To investigate the effect of block size on our inference, all analyses were repeated using shorter blocks (500bp; 9,085 unique bSFS configurations) which were obtained by dividing each 2kb block.

To facilitate comparison with previous studies, we fitted a series of increasingly complex models of divergence with gene flow (Fig. 4) to these data and estimated demographic parameters along with the genome-wide recombination rate *r* under each model. All demographic models included an instantaneous split at time *T*. We allowed effective population sizes *N*_*e*_ to differ between the two island populations and the ancestral population (M2 - M6). Additionally, we considered a model of divergence followed by exponential growth (or decline) in each population given by population specific growth rates *α* (M3). Asymmetric, bidirectional gene flow was modelled either as a continuous process occurring at a constant rate of *M* = 4*N*_*A*_*m* migrants per generation (M4 and M5), or as an instantaneous (bidirectional) admixture pulse affecting a fraction *f* of the admixted population (M6). We considered both an IM model with gene flow from time *T* to the present (M4) and a more complex history of isolation with initial migration (IIM) which assumes that migration ceases at time *T*_2_ (M5) [29]. To convert time estimates (scaled in 4*N*_*A*_ generations) into absolute time, we followed [21] and assumed a generation time of 20 years and a mutation rate *μ* = 2 × 10^−8^ bp^−1^ per generation.

**Fig 4.**
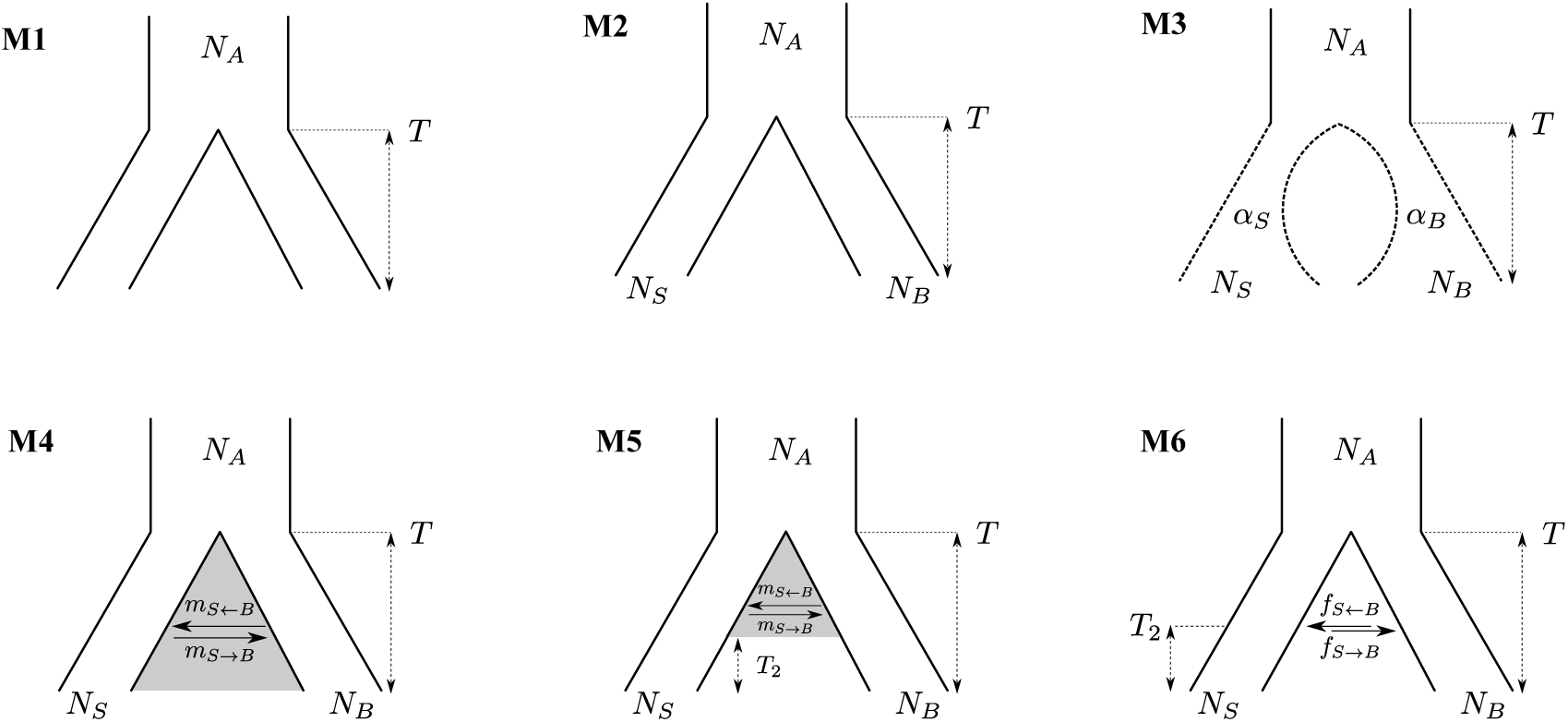
Models describing orangutan demography considered in this paper. All models describe a split between two populations at time *T* with an effective ancestral population size *N*_*A*_. In M2 - M6, the current population sizes are additional free parameters. A model of divergence followed by exponential growth (or decline) in each population is considered in M3. Continuous asymmetrical gene flow since the time of split is considered up to the present or with a stopping time *T*_2_ respectively in M4 and M5. Alternatively, an asymmetric *pulse* admixture at time *T*_2_ is considered in M6.

As expected, model support increased with increasing complexity for nested models (i.e. M1 vs. M2 and M4 vs. M5) (Fig. 5 and Table 1). The only exception was the IM model (M4) which did not increase support compared to a strict divergence history (M2). Interestingly, we found greater support for instantaneous admixture (M6) compared to a history of Isolation and Initial Migration (IIM) up to a time *T*_2_ (M5).

**Table 1.**
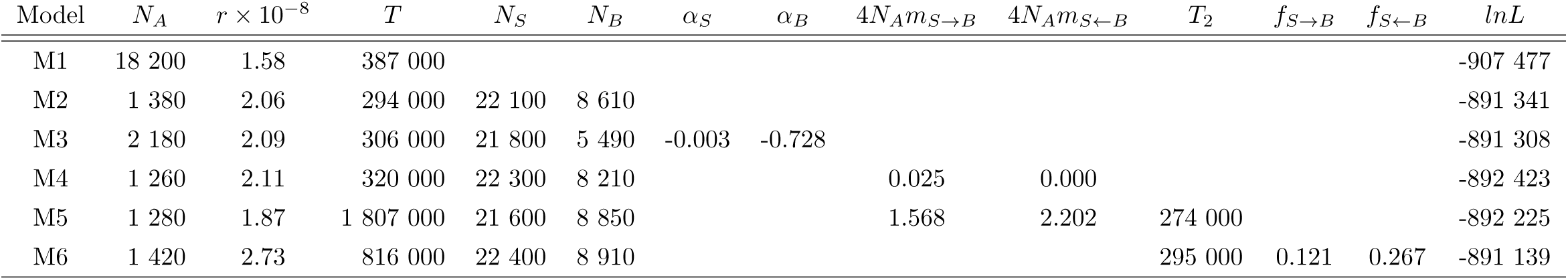
Point estimates for the demographic history of orangutan species obtained from 2kb blockwise data (*cf.* Fig. 4)

| Model | $N_A$ | $r \times 10^{-8}$ | $T$ | $N_S$ | $N_B$ | $\alpha_S$ | $\alpha_B$ | $4N_A m_{S \rightarrow B}$ | $4N_A m_{S \leftarrow B}$ | $T_2$ | $f_{S \rightarrow B}$ | $f_{S \leftarrow B}$ | $\ln L$ |
| --- | --- | --- | --- | --- | --- | --- | --- | --- | --- | --- | --- | --- | --- |
| M1 | 18 200 | 1.58 | 387 000 |  |  |  |  |  |  |  |  |  | -907 477 |
| M2 | 1 380 | 2.06 | 294 000 | 22 100 | 8 610 |  |  |  |  |  |  |  | -891 341 |
| M3 | 2 180 | 2.09 | 306 000 | 21 800 | 5 490 | -0.003 | -0.728 |  |  |  |  |  | -891 308 |
| M4 | 1 260 | 2.11 | 320 000 | 22 300 | 8 210 |  |  | 0.025 | 0.000 |  |  |  | -892 423 |
| M5 | 1 280 | 1.87 | 1 807 000 | 21 600 | 8 850 |  |  | 1.568 | 2.202 | 274 000 |  |  | -892 225 |
| M6 | 1 420 | 2.73 | 816 000 | 22 400 | 8 910 |  |  |  |  | 295 000 | 0.121 | 0.267 | -891 139 |

**Fig 5.**
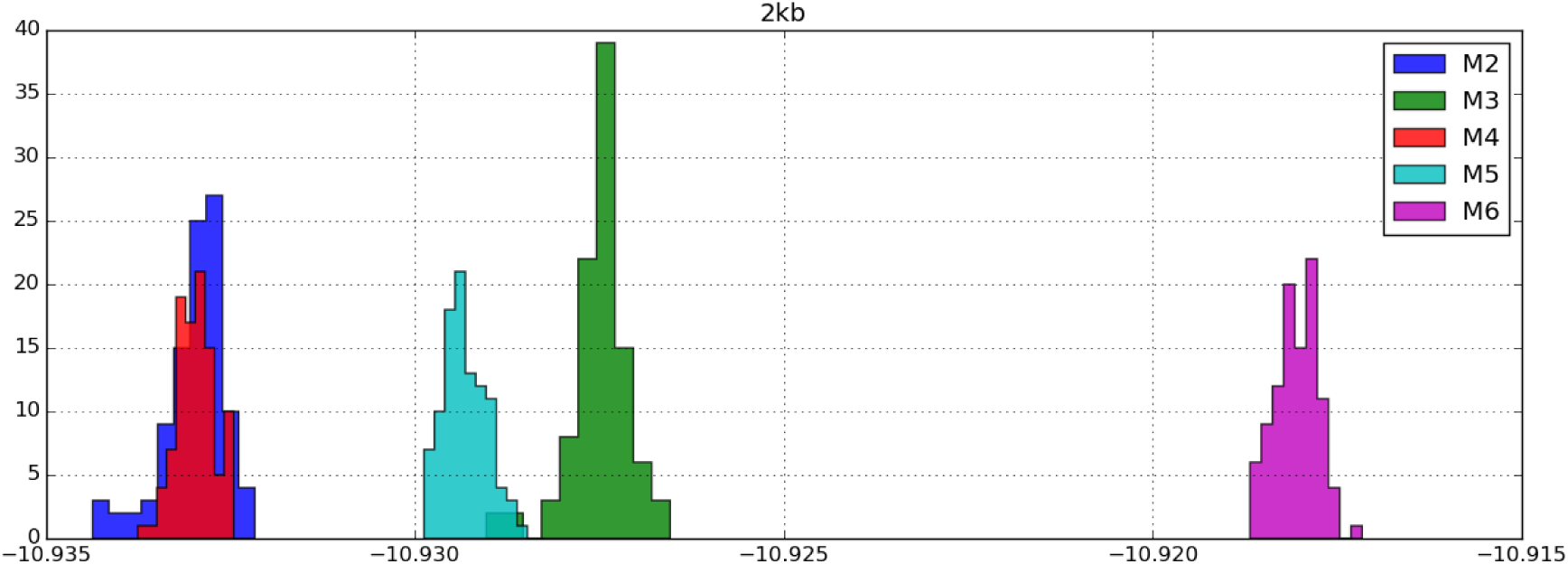
Relative fit of demographic models. Histograms of 100 evaluations of the Composite Likelihood (CL) using 1 million ARGs at the MCLE for each model (Table 1). The *x*-axis gives the *per block* CLs *i.e.* downscaled with respect to the number of blocks. Models further to the right fit relatively better than those to the left. Model M1 has the worst fit and is not shown (appears much further to the left)

Regardless of whether gene flow was modelled as a continuous process (M5) or a discrete admixture event (M6), our analyses reveal greater gene flow from Borneo into Sumatra than in the reverse direction. The maximum composite likelihood estimate (MCLE) under M6 (Table 1), the best supported model, suggests a much higher admixture fraction (*f*_*B*→*S*_ ≈ 0.2) and no significant admixture in the reverse direction (*f*_*S*→*B*_ ≈ 0.07).

Likewise, independent of any particular model, the effective size of the Sumatran species was 2.5 fold greater than that of the Bornean species. This is in agreement with previous studies [21] and mirrors the relative diversity in each species as measured by Watterson’s *θ* [30] (*θ*_*W*_ = 2.19 and 2.91 in 2kb blocks for the Bornean, Sumatran population respectively).

To determine the confidence in MCLE under M6, we carried out a full parametric bootstrap by simulating long stretches of sequence under the full ARG and determined 95 % Confidence Intervals (CI) as *±*2 SD (standard deviations) across bootstrap replicates (see Methods for details). The CIs in Table 2 (see also Fig. S2) indicate that we have relatively greater power to infer more recent aspects of orangutan history (*N*_*S*_, *N*_*B*_ and *T*_2_) compared to the time of initial divergence (*T*) and the size of the common ancestral population (*N*_*A*_). While the admixture fraction estimated from Sumatra to Borneo (*f*_*S*→*B*_) was not significantly different from 0, admixture estimates in the reverse direction had much tighter CI which clearly excluded zero.

**Table 2.** 95% confidence intervals obtained via a parametric bootstrap. 100 datasets were simulated given the point estimates of the 2kb analysis and model M6 (*cf.* Table 1). Bootstrap replicates were generated by cutting long (0.5 Mb) contiguous sequences into 2kb blocks. The confidence intervals were calculated as 2 standard deviations on either side of the maximum likelihood estimate.

| Parameter | MCLE ± 2SD |
| --- | --- |
| $N_A$ | 1,180 - 1,670 |
| $r \times 10^{-8}$ | 2.5 - 3 |
| $T$ | 695,000 - 936,000 |
| $N_S$ | 21,200 - 23,600 |
| $N_B$ | 8,400 - 9,420 |
| $T_2$ | 284,000 - 306,000 |
| $f_{S \rightarrow B}$ | 0 - 0.21 |
| $f_{S \leftarrow B}$ | 0.2 - 0.33 |

### Effect of block length and sample size

We assessed how block and sample size affects ABLE’s ability to infer two-population histories and recombination in two ways. First, we repeated the orangutan analyses using shorter blocks (500bp). Second, we used simulations to investigate how sampling additional genomes per population affects our inferential power.

### Block length

Comparing estimates based on 2kb blocks (Table 1) to shorter 500bp blocks (Table S1) suggest that most, but not all, aspects of the inference were fairly robust to block length. As expected, shorter blocks led to a greater uncertainty in model and parameter estimates (Table S2). Importantly however, even with 500bp blocks, M6 was identified as the best fitting model and we found broad overlap in 95 % CIs of parameter estimates with the 2kb analysis.

Both the divergence time *T* and the genome-wide recombination rate *r* were poorly estimated with 500bp blocks. While the 2kb analyses resulted in fairly stable inferences for *r* (≈2 *×* 10^−8^ bp^−1^ per generation) that agree with recombination estimates for humans [31], the 500bp estimates were 2-4 times greater and had very wide 95 % CIs (Table S2). However, note that the 95 % CIs of *T* under 2kb and 500bp do overlap for periods < 1 Mya.

To test whether our method has any inherent bias to overestimate recombination with shorter blocks, we simulated block-wise data under model M6 using the *r* estimates obtained from the 2kb data (Table 1). Applying ABLE to these simulated datasets and after taking into account the sampling strategy (*i.e.* M6 (2dp), Table S3), we noticed no significant overestimation of recombination rates. This may point to assumptions made by our approach on the type of recombination which may have been violated when considering 500bp blocks from the Orangutan data (see Discussion).

### Sample size

As expected, point estimates and power generally improved (Table S3 and Fig. S3) with increasing sample sizes. While some parameters, in particular *r*, appear non-identifiable with minimal sampling (a single diploid genome per species), all eight parameters of M6 are well estimated with just two or three diploid genomes. We observed a five fold improvement in accuracy for *r* and an up to two fold improvement for demographic parameters when increasing sampling effort from a single to two diploid genomes per population.

Perhaps surprisingly however, Fig. S3 suggests that for histories similar to that inferred for the two orangutan species, we can expect at best slight improvements in power when adding a third diploid genome per population. Given that analysing three diploid samples per population almost triples the computation time (Fig. S4), this suggests that (at least in the case of orangutans) analysing a total of four diploid genomes is a good compromise between information and computational cost.

### Model misspecification

When analysing real data, the underlying true demography is of course unknown. Thus, an important question is to what extend alternative demographic histories can be distinguished. We evaluated the ability of ABLE to distinguish between three progressively nested models (M1, M2 & M6, see Fig. 4). For each scenarios we simulated 20 datasets (see Table S4) and compared the overall fit to the true and alternative models. As expected (given that models were nested) data generated under simple models did not give a better fit to more complex histories (Fig. 6). In contrast, data generated under more complex histories showed a worse fit to simpler scenarios than the truth.

**Fig 6.**
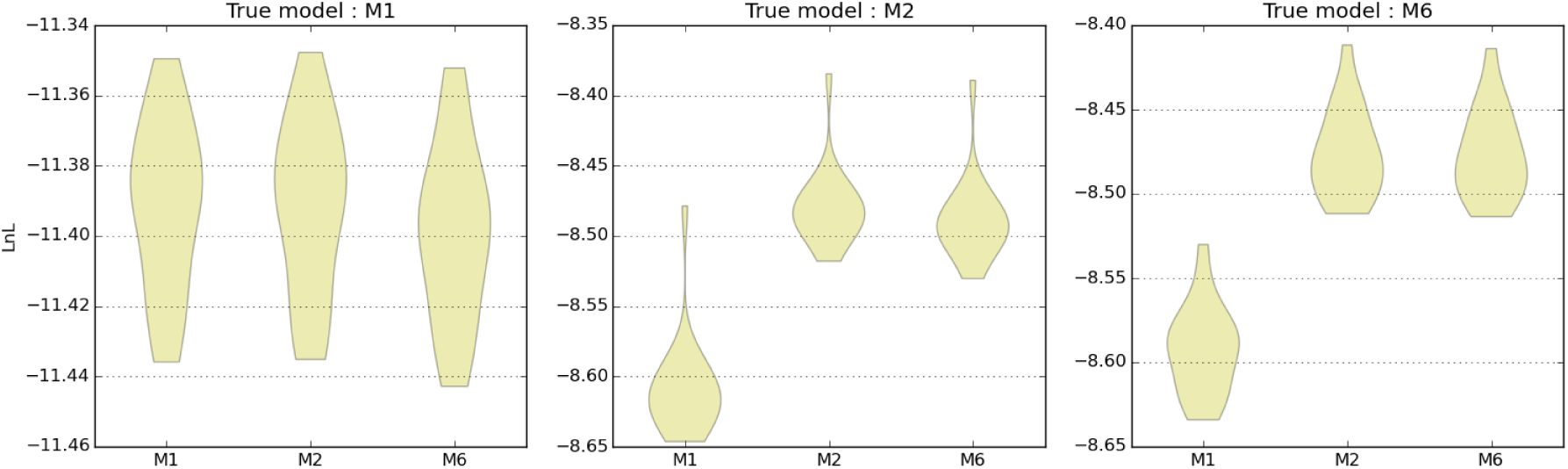
Nested model fit under the true and alternative models. Point LnLs were evaluated at the MCLE under the true and alternative models and for the 20 simulated datasets/model. Higher LnL values (*i.e.* closer to zero) indicate a better fit to the simulated data.

However, given the increasing dimensionality of the likelihood surface for more complex models, the similar LnL values for nested models did not imply that the actual ML estimates of demographic parameters under the simpler models were a subset of the corresponding estimates under the more complex models (see Fig. S5, Fig. S6 and Fig. S7). For instance, given M1 as the true model (Fig. S5), the population split time was largely overestimated under M6 as this model also consists of another confounding demographic feature, a pulsed admixture event subsequent to divergence. Interestingly, the genome-wide recombination rate was fairly consistently estimated among the various models while the ancestral population size was consistently underestimated.

To further investigate the ability to correctly identify complex demographies involving post-divergence admixture, we generated 20 simulated datasets under the most complex model considered in the orangutan analysis (M6). We considered 9 different divergence/admixture times which varied from 900-150Kya and 600-75Kya respectively with all other values being kept constant (Table S4) and compared LnL at the true parameter values with MCLE estimated for lower dimensional models M1 & M2. Point LnLs were also calculated for variants of the M1 & M2 models (M1R & M2R respectively) where the true divergence time was instead replaced by the true admixture time of the simulated dataset (Fig. S8). This analysis illustrates that the ability to identify population divergence and subsequent admixture depends crucially on the interval between these events. When the interval is approximately *T − T*_2_ < 0.3 coalescent units, M2R and M6 become indistinguishable (Fig. S8), which explains the difficulty of distinguishing between M2 and M6 (Fig. 6).

### Comparison between ABLE and ∂a∂i

Using simulated datasets, we compared the bias and accuracy of ABLE to that of a popular SFS-based method ∂a∂i [3]. We simulated data under three progressively nested models M1, M2 & M6 (Fig. 4. For each model, a total of 10 datasets were simulated (see Table S4). Each dataset consisted of 5 diploid genomes/population. The ∂a∂i analyses were based on the whole sample, while ABLE was only used on subsamples of two diploid genomes per population.

Despite the fact that ABLE used less than half of the data, it performed as well and in some cases better as ∂a∂i (Fig. S9, Fig. S10 and Fig. S11). While both methods underestimated ancestral population size (*N*_*A*_) for models M1 & M2 and overestimated the admixture fraction under M6, ABLE in general gave less biased estimates of divergence and admixture times than ∂a∂i.

## Discussion

We have developed a flexible, efficient and widely applicable simulation-based approach to simultaneously infer complex demographic histories and average genome-wide recombination rates under the full ARG. This method overcomes the limitations of previous approaches that either ignore recombination [3,4], use fixed estimates [19], approximate recombination as a Markov process along the genome [8,11–13] or are limited by the type of population histories they infer [32,33]. Using the bSFS as a data summary, ABLE captures linkage information at the scale of hundreds to thousands of base pairs across the genome, and allows researchers to efficiently fit realistic demographic models across the variety of genome scale data sets that are becoming available for a rapidly growing number of species.

The quick asymptotic convergence of the bSFS approximated by ABLE to the expected bSFS under various demographic scenarios (Fig. 3) in the absence of recombination is reassuring and distinguishes our method from related multi-locus approaches that integrate over possible genealogies locus by locus [19]. Moreover, the fit of the expected bSFS to data simulated with long range linkage is encouraging (Fig. S12) highlights the potential of the bSFS for the purpose of demographic inference and beyond.

### Orangutan history

The best fitting demographic model (M6) suggests that the two *Pongo* species diverged 650 - 1000 kya and experienced a burst of admixture around 300 kya. Given the Pleistocene history of periodic sea-level changes in South East Asia [24], such a scenario of secondary contact seems biogeographically more plausible than continued migration. Interestingly, our estimates of the divergence time under M6 are consistent with previous estimates based on the SMC [8,22] and also agree well with species splits estimated for other island-endemic mammals in SE Asia [24].

Overall, our results are in general agreement with previous analyses regarding the absence of recent gene flow (< 250 kya) between Bornean and Sumatran orangutans (see Tab. S21-1 & S21-2 in [21]). Likewise, our inference of a larger *N*_*e*_ in Sumatran compared to Bornean orangutans agrees with relative measures of nucleotide diversity and previous analyses using various types of data [12,19,21,23]. While we infer a contraction for the Bornean population under M3, in agreement with the simpler models explored by [23], sampling at finer spatial scales would be required to resolve the issue of substructure in both the Sumatran and Bornean populations.

Reassuringly, the time of secondary admixture under M6 agrees with the estimated split time between the two *Pongo* species for simpler models M1-M4 (Table 1) which are similar to those considered by Locke *et. al.* [21]. Using the joint SFS (*δaδi*, [3]), Locke *et. al.* [21] estimate a species divergence time of 400 kya, which is somewhat older than our estimate (250-300 kya) under M1-M4. However, a similar difference in estimates has already been noted by the Hidden Markov Model approach of Mailund *et. al.* (see Supplemental Text S2, [13]) which models a simplified demography of speciation with continuous gene flow and recombination using whole genome data.

### Absolute model fit and the effect of selection

Like most demographic inference methods, ABLE assumes selective neutrality. Furthermore, efficient calculation or approximation of the bSFS relies on the assumption that blocks are statistically exchangeable, and so ignores heterogeneity in mutation and recombination rates.

We can visualize the absolute fit of our demographic model to the data by comparing the observed distribution of bSFS configurations to that expected under M6 (obtained using 50 million simulated blockwise ARGs). If the data were generated entirely by the inferred demographic history alone, we would expect the most common bSFS configurations to fit this expectation most closely (see Fig. S12). In contrast, Fig. 7 shows that, irrespective of which demographic model we assume, some aspects of the data are poorly captured. In particular, the most likely bSFS configurations are over-represented in the data. This effect is strongest for configurations with few (or no) mutations (shown in blue), which is compatible with background selection [34] and/or positive selection affecting a fraction of blocks.

**Fig 7.**
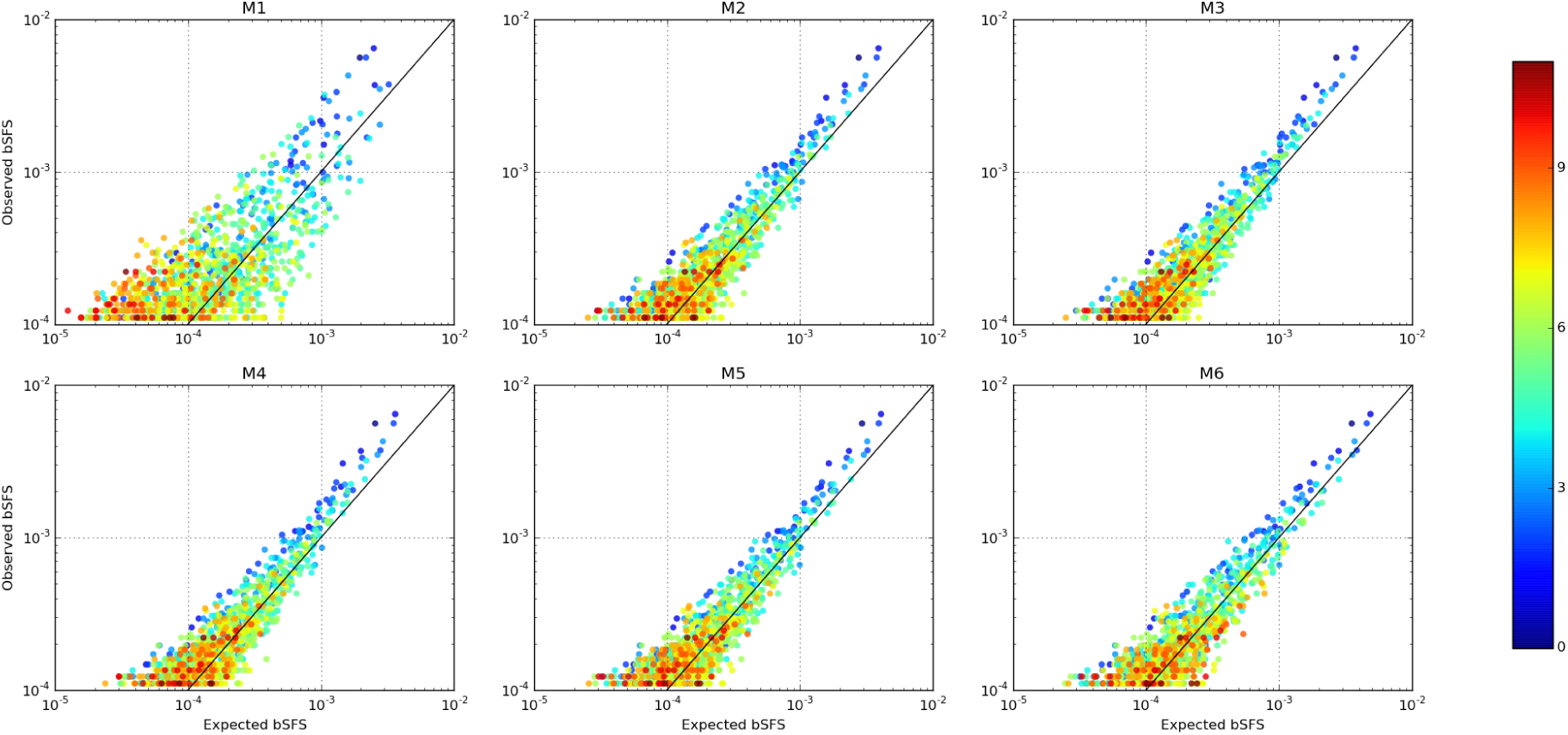
Absolute model fit to the observed 2kb bSFS for the most common configurations. Each point here represents a unique mutational configuration making up the bSFS. The expected bSFS (*x*-axis) was generated with ABLE using 50 million ARG’s at the MCLE for each model (Table 1) and plotted against the observed bSFS (*y*-axis) from the orangutan data. The diagonal black line indicates the perfect match between the expected and observed. The colours represent the total number of SNPs contained in each configuration.

If background selection at linked sites is indeed responsible and a fraction of selective targets is weakly deleterious, we would expect increasingly biased *N*_*e*_ estimates going backwards in time. This could explain why we estimated a much smaller effective size for the ancestral population (Table 1 and Table S1) than previous studies [12,19,21,23], while our *N*_*e*_ estimates for the two current populations agree fairly well [13]. As expected, this *signature* of linked selection disappears as we consider a bSFS with reduced block size (Fig. S13). Going further, it will be interesting to explore the possibility of jointly inferring demography and various forms of selection using the bSFS [35].

### Effect of block length and sample size

An interesting property of the bSFS is that it collapses to the SFS in both the limits of minimal block length (one base) and maximal block length (all data in a single block). At both extremes, all linkage information is lost and so the information contained in the distribution of bSFS types must be optimal at an intermediate scale. A sensible starting point to finding an informative block length for a particular dataset is to ask at what length the number of unique bSFS configurations is maximised (Fig. S14). Crucially however, the most informative block length will depend on *μ/r* and the demographic history and parameters in question. For example, a burst of recent admixture generates an excess of long blocks with excessive LD. Thus, an interesting direction for future work would be to integrate CL estimates based on the bSFS over a range of block sizes which should improve the power to infer recent demographic events.

Our finding of larger *r* estimates when using shorter blocks for the Orangtuan data was surprising. Given that our method ignores heterogeneity in both *r* and *μ*, both of which increase auto-correlation across short distances, we expected to find the opposite, *i.e.* a decrease in *r* estimates for shorter blocks. However, our simulation analysis showed that ABLE gives relatively unbiased estimates of *r* for short (500bp) blocks when inference was performed using 2 or more diploid samples per population (Table S3). A plausible explanation for the larger estimates in Orangutan data could be gene conversion as it could potentially inflate *r* at short blocks with a diminishing effect on the bSFS for blocks that are longer than the typical conversion tract length of several hundred bases (see Table 2 in [36]). It should be possible to use this dependence on block length to develop explicit estimators for gene conversion and cross-over rates.

Even under a complex demography such as M6, our simulation-based power analyses indicate that most demographic parameters can be reasonably recovered with only a single diploid genome per population (Table S3). Increasing sample size to two diploid genomes more than halved the standard deviation in estimates for some parameters, most notably the recombination rate (Fig. S3). However, a further increase in sample size gave a negligible improvement, despite the considerable computational cost (Fig. S4) involved: the number of unique bSFS configurations increased more than three fold with three rather than two diploid genomes per population. This diminishing return with increasing sample size (in terms of sequences), is a fundamental property of the coalescent [37,38]: going backwards in time, larger samples in each species are likely to have coalesced down to a small number of lineages (see Fig. 3 in [38]) before the admixture event and so could not contribute much additional information about older demographic processes.

### The SFS and the bSFS

In this paper, we have explored the intuition that accommodating for linkage via the bSFS is likely to produce better inference results than those based solely on the SFS as the latter is only a function of the expected length of genealogical branches [5,6]. Previously it has been shown that in a sample from a population having undergone a single instantaneous bottleneck, the bSFS from fewer individuals (*n* = 5) contains significantly more information on the historical event compared to the SFS using *n* = 20 (see Fig. 3 in [20]). Likewise, our analysis comparing ABLE with SFS-based ∂a∂i [3] for progressively complex subdivided population scenarios (M1, M2 & M6) resulted mostly in similar inferences of the population history and with a slight improvement for admixture rates when using ABLE (Fig. S9, Fig. S10 and Fig. S11). Of note however, we only make use of a random subset (2 diploid genomes for the ABLE analysis) of the simulated data (5 diploid genomes for the ∂a∂i analysis) under each scenario. This *increase* in performance can be explained by the fact that the bSFS captures a wide range of historical signatures within all its blockwise mutational configurations which greatly surpass those considered by the SFS [18,20]. However, this significant increase in mutational configurations [18] also implies an increase in the cost of computing the bSFS (Fig. S4) and it may be fruitful to narrow down parameter space using SFS-based approaches such as ∂a∂i [3] prior to an ABLE analysis.

### Limits to inference

While our choice of models was guided by previous knowledge of the demographic history of orangutans [13,21–23], it remains to be determined what the limits of model complexity and identifiability are with our approach and to what degree the distribution of bSFS patterns overcomes the non-identifiability of the SFS [7,39,40]. Unlike analytic likelihood calculations (*e.g.* [18]), there is no significant increase in computational cost with increasing model complexity when approximating the likelihood for a given point in parameter space. However, searching parameter space carries an obvious and rapidly increasing cost with greater model complexity. Like all approximate likelihood approaches, ABLE requires the user to make careful choices about the number of parameters, the number of genealogies to sample per point in parameter space, the search bounds for the MCLE all of which are crucial elements of the optimization strategy [4]. In this regard, we suggest that simple pilot analyses varying some or all of the factors mentioned above (see Fig. S15 and Fig. S14) should help to inform the inference strategy.

It is also clear that, independent of the inference approach, the information in the data is finite so there must be a hard limit on how realistic a history one can hope to infer. Thus, the fact that ABLE can, in principle, be used for fitting any demographic model puts the onus of constraining inference to scenarios that are both statistically identifiable and biologically interpretable on the user. Evaluating the relative fit of simpler nested models is an important sanity check on the limits of information in the data. For instance, comparing relative model fits between 2kb and 500 bp datasets (Fig. 5 and Fig. S16 respectively) highlights the apparent limits of using a bSFS with the latter block size in discriminating between models.

The inferential approach implemented in ABLE makes use of the coalescent simulator ***ms*** [41] for sampling blockwise genealogies or ARGs. In principle ABLE can accommodate other simulators and is thus amenable to include additional processes such as linked selection [42,43]. Another interesting avenue for further research is to apply approximate composite likelihoods based on the bSFS in sliding windows along the genome. Such an approach would not only help improve upon recombination maps for non-model organisms but could also provide a robust framework to identify *outlier regions* of the genome under positive selection and/or affected by introgression from another species.

## Methods

### Approximating the bSFS

#### A single population

It is easiest to first consider the simpler case of non-recombining blocks and a sample of *b* genomes from a single panmictic population. We assume an arbitrary population history which is entirely described by a vector of parameters Θ In the simplest case, Θ consists of the scaled mutation rate *θ*= 4*N*_*e*_*μ*, where *N*_*e*_ is the effective population size and *μ* the mutation rate per site per generation.

The branches of a given genealogy corresponding to our population sample can be partitioned into a vector <u>*t*</u> whose entries *t*_*i*_ *∈* [*t*_1_*, t*_2_*, …, t*_*b*−1_] denote the total length of all branches with *i* descendants (Fig. S1). The probability of observing *k*_*i*_ mutations on a branch class *t*_*i*_ is given by a Poisson distribution with rate parameter *θt*_*i*_ *>* 0:

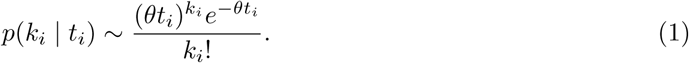

Because mutations occur independently on different branch types, the joint probability of seeing a specific configuration <u>*k*</u>_*j*_ = {*k*_1*,j*_*, k*_2*,j*_*, …, k*_*b-*1*,j*_} in a sequence block *j* and for a given branch length vector <u>*t*</u> is then a product of Poisson distributions

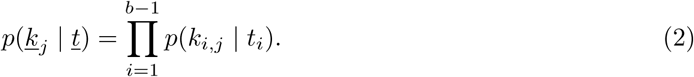

The likelihood *ℒ*(Θ) at a point in parameter space Θ ∈ ℝ^+^ is calculated as

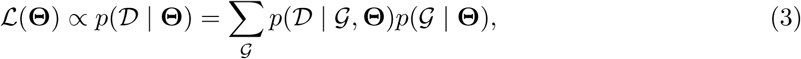

where *𝒢* is the (unknown) genealogy and *𝒟* the data [44]. Summarizing genealogies *𝒢* by <u>*t*</u> and *𝒟* by <u>*k*</u>_*j*_ and drawing *ℳ* random samples of <u>*t*</u> from *p*(*<u>t</u> |*Θ), the Monte Carlo approximation of Eq. 3 can be obtained

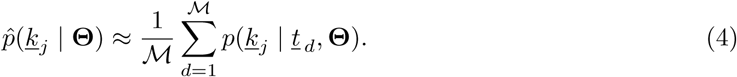

In theory, each block in a dataset might have a unique bSFS configuration. In practice however, for short blocks spanning a handful of SNPs (*e.g.* < 10), the number of observed bSFS configurations will be much smaller than the number of blocks. Assuming that blocks are equivalent and independent, that is, they have the same length, per base mutation and recombination rates and are unlinked, we can summarize the entire genome into *blockwise data* (Fig. 1) by counting the number of each unique bSFS type 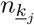. Thus, the approximate joint composite log likelihood for a sample of *n* genomes is given as

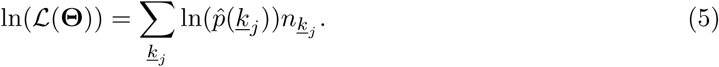

#### Multiple populations

The Monte Carlo approximation detailed above extends itself easily to the joint bSFS [6,18] for multiple populations. Assuming a sample from *X* populations, the (unfolded) joint bSFS defines 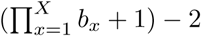 site types, where *b*_*x*_ denotes the number of genomes sampled from population *x*. Some branches will be specific to a single population while others are shared between populations. Thus the vectors <u>*t*</u> and <u>*k*</u> have entries corresponding to the joint bSFS. Note that one specific configuration which we denote as *k*_0_ refers to monomorphic blocks.

#### The Ancestral Recombination Graph

In the presence of recombination, the ancestry of a sequence block is described by the *ancestral recombination graph* 𝒜 [1] which can be partitioned into a set of marginal genealogies corresponding to the non-recombining segments that make up the block [45]. Here, Θ consists of the scaled mutation rate *θ* and the scaled recombination rate *ρ* =4*N*_*e*_*r*, where *r* is the recombination rate per site per generation. For a given *A*, let *S* be the number of non-recombining blocks with respective (sequence) lengths *w*_1_*, w*_2_*, …, w*_*S*_ such that the size of the sequence block 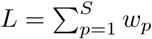. Let <u>*t*</u>_*p*_ be the marginal branch length vector for each non-recombining segment *p*. The total length of the *i*^*th*^ branch class over the graph 𝒜 is then given by

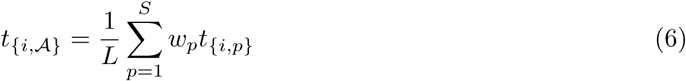

Following Eq. 1, we can write the joint probability of observing a specific bSFS configuration over the entire recombining block as *p*(*k*_*{i,𝒜}*_ *| t*_*{i,𝒜}*_) ∼ *Poisson*(*k*_*{i,𝒜}*_; *θt*_*{i,𝒜}*_) (analogous to Eq. 2). Drawing *M* random samples of *A* from *p*(𝒜 *|*Θ) and replacing *p*(*k*_*j*_ *| t_d_*, Θ) with *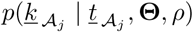* in Eqs. 4 and 5 gives the approximate likelihood for a point in parameter space Θ, ρ∈ℝ^2+^ (see also [19]). However note that Θ ρ∈ℝ^2+^ can be too restrictive a criterion for some parameters of complex demographies such as coefficients of exponential population expansion/contraction where α _*S*_, α _*B*_ ∈ℝ (see Fig. 4).

### Implementation

The ABLE implementation includes a seamless integration (invisible to the user) of the simulator *ms* [41] for sampling genealogies from *p*(𝒢 *|* Θ) or *p*(𝒜 *|* Θ). Crucially, for each simulated genealogy, we only record the total branch lengths of all SFS classes *t*_*{i,𝒜}*_ in each ARG. This is a sum over marginal genealogies contributing to the ARG, each weighted by its length. From these we can tabulate the probabilities (conditional on *𝒢*) of all bSFS patterns compatible with that ARG. This task is extremely efficient compared to previous *multi-locus* methods that sample *𝒢* separately for each locus (see [19,46]).

Note that ABLE differs from previous, analytic calculations based on the distribution bSFS configurations in an important way. Lohse *et al.* [18] tabulate probabilities of all bSFS configurations up to a maximum number of mutations (*k*_*max*_) in each category and lump all configurations *> k*_*max*_ mutations.

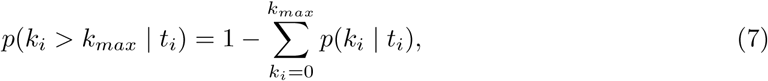

and

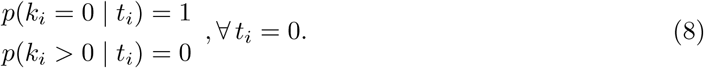

Bounding the table of mutational configuration in this way makes analytic computations feasible and ensures that the table of probabilities sums to unity. However, choosing *k*_*max*_ involves a trade-off between computational efficiency (low *k*_*max*_) and information (high *k*_*max*_). In contrast, ABLE only computes probabilities for mutational configurations that are observed in the data without setting any bounds on the space of possible configurations.

ABLE is implemented in C/C++, follows closely the command-line structure of *ms* [41] along with a brief configuration file with additional instructions and is freely available for download from https://github.com/champost/ABLE.

### Data processing

Raw reads were downloaded from the NCBI Short Read Archive (SRA) for two individual genomes each from Borneo (B) and Sumatra (S): KB5405 (B, male, SRS009466), KB4204 (B, male, SRS009464), KB9258 (S, male, SRS009469), KB4361 (S, female, SRS009471). Mean Depth of Coverage was between 7.25 and 8.06 per individual. The alignment was performed using BWA-MEM (arXiv:1303.3997), with a re-alignment step using GATK (PMC2928508). For each sample we estimated a 95 % Depth of Coverage interval using BEDTools [47]. To call genotypes we used a simple approach that was previously used to infer the genotype of species distantly that were aligned to a distant reference genome (PMC4053821; PMC4284605). To do so, we generated pileup files using samtools ‘mpileup’ (0.1.19) (PMC2723002) with default settings. Pileup files were then filtered, for each sample, using the following criteria:

- Minimum Depth of Coverage *>*= 4 reads with mapping quality *>*= 30
- Excluded all sites in region of high DoC (top 5%) (coded as N to avoid copy number variant)
- Excluded all sites within 5bp of an indel (coded as N to avoid indel missalignments)
- Only bases with quality *>*= 30 within reads with mapping quality *>*= 30 were used.
- Minimum fraction of reads supporting heterozygous (variant allele frequency [VAF] *>*= 0.3). All sites that did not pass this criteria (0 *<* VAF *<* 0.3) were coded as missing (N).

We also masked any non reference allele that were within 3bp of an indel and checked for allele balance at heterozygous calls (e.g. each allele was covered by *>* 20% of the reads at that position). Thereafter, we binned the genome into non-overlapping blocks of fixed length *l* = 2*kb* and sampled the first 0.8 *× l* = 1600 bases in each block that passing filtering in all individuals (a python script is available upon request). Blocks with fewer bases post filtering were excluded. The 500bp dataset was generated by partitioning each post-filtered 1.6kb block into four blocks of equal size. The 500bp and 2kb block datasets used in this study (in bSFS format and only genotypes) are available for download from the aforementioned website.

### Optimization

Because ABLE approximates the likelihood function (Eq. 5) using Monte Carlo simulations – which induces some variability in the CL obtained (Fig. S15) – algorithms based on the gradient of the CL surface (*e.g.* [3,9]) are not reliable [4]. In addition, due to the possibility of multiple local optima in the likelihood surface, we adopted a two-step search heuristic.

We initially searched parameter space between broad, user-specified non-linear bounds as part of a *global search* step. Search bounds during this step spanned several orders of magnitude for all parameters. Upper bounds of some parameters were set on the basis of simple data summaries: *e.g.* effective population sizes were bounded by Watterson’s _*W*_ [30]. 50,000 ARGs were used to approximate the CL at each point in 10 replicate global searches. These were then used to set narrower bounds for a *local search* based on 500,000 ARGs/point which was repeated 20 times. In Table 1 and Table S1, we report the best MCLEs whose likelihoods have been evaluated using 1M ARGs. For some models for which replicate local searches did not converge sufficiently, a second round of local searches was used.

ABLE employs several search algorithms implemented in the Non-Linear optimization library (NLopt version 2.4.2, [48]). Both global and local searches used the improved penalization provided by the Augmented Lagrangian algorithm [49] to navigate the non-linear delimitation of parameter space. A Controlled Random Search with rules for the evolution of a population of points given by the Local Mutation algorithm [50] was used for global searches. Local searches used the Subplex algorithm [51], a variant of the Nelder-Mead simplex with start points that were randomly chosen within the parameter bounds set by the global searches.

Finally, tolerances for terminating MCLE searches were determined by probing the CL surface (*e.g.* Fig. S15). The command lines and configurations used to analyse the Orangutan data are available as part of the supplement (see Text. S1, Text. S2, Text. S3, Text. S4, Text. S5, Text. S6 and Text. S7).

### Parametric bootstrap and simulation analysis

While the CL is a statistically consistent estimator of demographic parameters and recombination (in the limit of large data, [52]), it suffers from severe overconfidence because correlations between blocks due to their physical linkage are ignored. To obtain meaningful measures of confidence, we conducted a full parametric bootstrap under the best fitting model (M6) and parameter estimates (Table 1). We simulated 100 replicate datasets of 164 Mb each using a modified version of *ms* [41] and under the best model (*i.e.* M6) and MCLE (Table 1 and Table S1). Blocks in each dataset were assumed to be completely linked (given our estimate of per site *r*) across 0.5 Mb stretches of sequence. These simulations represent an extreme case of linkage and are thus conservative. Indeed our real data contain large gaps between blocks especially due to the highly repetitive nature of the orangutan genome. As we wish to know the *local variability* of the bootstrap inferences around the MCLE obtained from the orangutan data, we only carried out local searches for each bootstrap replicate (using the boundaries and step sizes obtained in the analysis of real data, see above).

The simulation based power test exploring the effect of sample size (1 - 3 diploid genomes per population) was based on inferences using simulated data followed by a full parametric bootstrap. Given the computational effort required (see Fig. S4), we restricted our study to 500bp blocks with values for the demographic parameters chosen to represent the results inferred from the real data under M6 (Table 1 and Table S1). Parametric bootstrap datasets were generated with linkage (under the full ARG) exactly analogous to the bootstrap in the real data analysis.

### Evaluating nested model misspecification

To evaluate model misspecification we compared the overall fit of several models to a dataset simulated under a specific model. Thus, datasets were generated under three 2 population demographic scenarios M1, M2 & M6 (4). For each scenario, 2 diploid genomes/population sample were simulated (using *ms* [41]) each of which had a size of 200 Mb (made up of independent 1 Mb blocks) and 2 diploid samples/population. The values used for the simulation can be found in Table S4. Under any given model, the MCLE search strategy consisted of a three global searches of the parameters bounds and finally a local search. The final likelihoods were evaluated using 1M genealogies.

Assuming that the parameter values used to simulate data would have been close to the inferred global maximum under both the true/alternative models, we also attempted an illustration of model choice with ABLE by comparing LnLs under M1, M2 and M6 in the tricky situation when M6 is the true model (Fig. S8). 20 datasets were simulated under model M6 and 9 different split/admixture times. Split times varied from 900-150Kya whereas admixture times varied from 600-75Kya and sample sizes were the same as in the previous section. Model fit was assessed using point LnLs calculated at the true parameter values of each simulated dataset which meant using only a subset of those values for the lower dimensional models M1 & M2. Point LnLs were also calculated for variants of the M1 & M2 models (M1R & M2R respectively) where the true split time was instead replaced by the true admixture time of the simulated dataset.

### Comparison between ∂a∂i and ABLE

We compared the performance in terms of parameter inference between ∂a∂i [3] and ABLE when the latter uses a subset of every simulated dataset under M1, M2 & M6. Akin to the previous section, we simulated 2 population demographic scenarios under M1, M2 & M6. Each simulation consisted of 5 diploid genomes/population sample and each genome was made up of 1M 2 Kb blocks (*i.e.* 200 Mb in size). A total of 10 datasets were simulated for each of the three scenarios (see Table S4). Each dataset was either summarized as the folded SFS (for a subsequent ∂a∂i analysis) and by the folded bSFS by randomly sampling 2 diploid genomes from each population (for a subsequent ABLE analysis).

Parameter inference under M1 & M2 for both ∂a∂i and ABLE analyses was performed using 10 independent (local) searches on each simulated dataset. For the M6 scenario, ABLE analyses followed a global search with successive refinement due to the high dimensional search space while ∂a∂i analyses were consistent with its previous strategy. Python scripts defining the models M1, M2 & M6 (see Text. S8, Text. S9 and Text. S10) to facilitate a ∂a∂i analysis have been made available as part of the supplement to this paper.

## Acknowledgments

We thank Stuart Baird, Lynsey Bunnefeld, Brian Charlesworth and Graham Stone for helpful discussions throughout this project. KL was supported by an Independent Research fellowship from the Natural Environment Research Council (NE/L011522/1). CBR and MJH were supported by grants from FAPESP (BIOTA, 2013/50297-0 to M.J.H. and Ana Carnaval), NASA through the Dimensions of Biodiversity Program, and National Science Foundation (DOB 1343578 and DEB-1253710 to MJH). LAFF was supported by a Junior Research fellowship from Wolfson College (University of Oxford) and a European Research Council grant (ERC-2013-StG-337574-UNDEAD). This research was supported by National Science Foundation Grants CNS-0958379, CNS-0855217, ACI-1126113 to the City University of New York High Performance Computing Center at the College of Staten Island. We also wish to thank Silicon Mechanics and their Research Cluster Grant program for the donation of the high-performance computing cluster that was used in support of this research.

## Supporting Information

### Tables

**Table S1.** Point estimates for the demographic history of orangutan species obtained from 500bp blockwise data (*cf.* Fig. 4).

| Model | $N_A$ | $r \times 10^{-8}$ | $T$ | $N_S$ | $N_B$ | $\alpha_S$ | $\alpha_B$ | $4N_A m_{S \rightarrow B}$ | $4N_A m_{S \leftarrow B}$ | $T_2$ | $f_{S \rightarrow B}$ | $f_{S \leftarrow B}$ | $\ln L$ |
| --- | --- | --- | --- | --- | --- | --- | --- | --- | --- | --- | --- | --- | --- |
| M1 | 18 000 | 3.34 | 413 000 |  |  |  |  |  |  |  |  |  | -1 653 152 |
| M2 | 1 640 | 4.68 | 314 000 | 21 900 | 9 090 |  |  |  |  |  |  |  | -1 632 879 |
| M3 | 1 970 | 4.39 | 347 000 | 23 000 | 4 120 | 0.019 | -1.132 |  |  |  |  |  | -1 632 852 |
| M4 | 1 920 | 5.81 | 317 000 | 21 900 | 8 020 |  |  | 0.027 | 0.000 |  |  |  | -1 633 028 |
| M5 | 1 800 | 5.13 | 2 800 000 | 21 200 | 8 530 |  |  | 4.961 | 6.148 | 289 000 |  |  | -1 631 767 |
| M6 | 1 480 | 8.67 | 1 345 000 | 22 500 | 9 500 |  |  |  |  | 329 000 | 0.010 | 0.167 | -1 628 548 |

**Table S2.** 95% confidence intervals obtained via a parametric bootstrap. 100 datasets were simulated given the point estimates of the 500bp analysis and model M6 (Table S1). Bootstrap replicates were generated by cutting long (0.5 Mb) contiguous sequences into 500bp blocks. The confidence intervals were calculated as 2 standard deviations on either side of the maximum likelihood estimate.

| Parameter | MCLE ± 2SD |
| --- | --- |
| $N_A$ | 1,160 - 1,810 |
| $r \times 10^8$ | 5.8 - 11.5 |
| $T$ | 822,000 - 1,869,000 |
| $N_S$ | 19,400 - 25,700 |
| $N_B$ | 8,100 - 10,890 |
| $T_2$ | 285,000 - 374,000 |
| $f_{S \rightarrow B}$ | 0 - 0.06 |
| $f_{S \leftarrow B}$ | 0.08 - 0.26 |

**Table S3.** Point estimates under M6 (*cf.* Fig. 4) using 500bp blocks and 1/2/3 diploid genomes per population respectively. The MCLE for each sampling scheme is shown in gray, the true values used for the simulations in white.

| Model | $N_A$ | $r \times 10^{-8}$ | $T$ | $N_S$ | $N_B$ | $T_2$ | $f_{S \rightarrow B}$ | $f_{S \leftarrow B}$ | $\ln L$ |
| --- | --- | --- | --- | --- | --- | --- | --- | --- | --- |
| M6 | 1 250 | 2.50 | 1 000 000 | 16 250 | 8 750 | 500 000 | 0.200 | 0.500 |  |
| M6 (1dp) | 1 352 | 12.2 | 1 732 345 | 13 343 | 8 404 | 439 427 | 0.389 | 0.295 | -3.314791 |
| M6 (2dp) | 1 588 | 2.89 | 1 080 289 | 16 360 | 9 055 | 502 330 | 0.283 | 0.426 | -5.531729 |
| M6 (3dp) | 1 857 | 2.77 | 1 095 628 | 15 655 | 8 598 | 492 806 | 0.177 | 0.446 | -7.040805 |

**Table S4.**
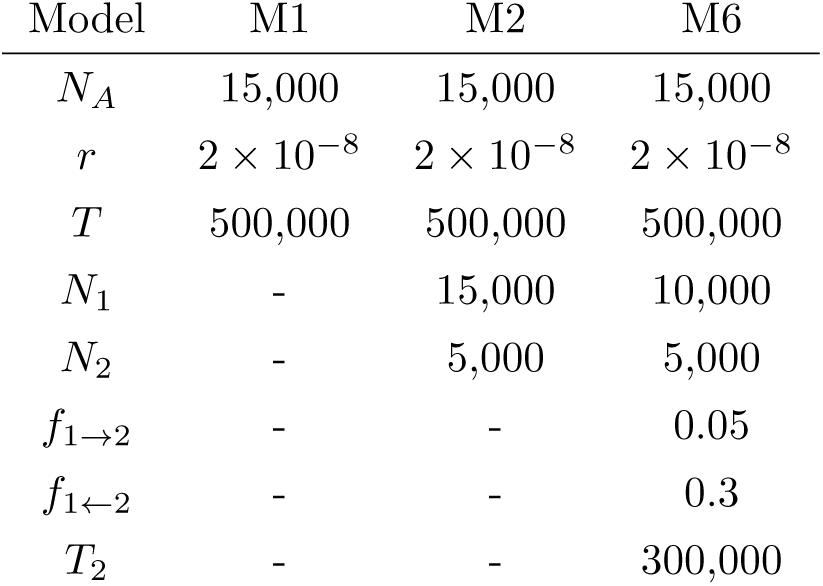
True values for generating datasets under the M1, M2 and M6 models. Simulations were performed with the values specified below for studying model misspecification under progressively nested models (*cf.* Fig. 4). *r* stands for the recombination rate/base pair/generation and we assumed 20 years/generation. All event times (*T* and *T*_2_) have been specified in years.

### Figures

**Fig. S1.**
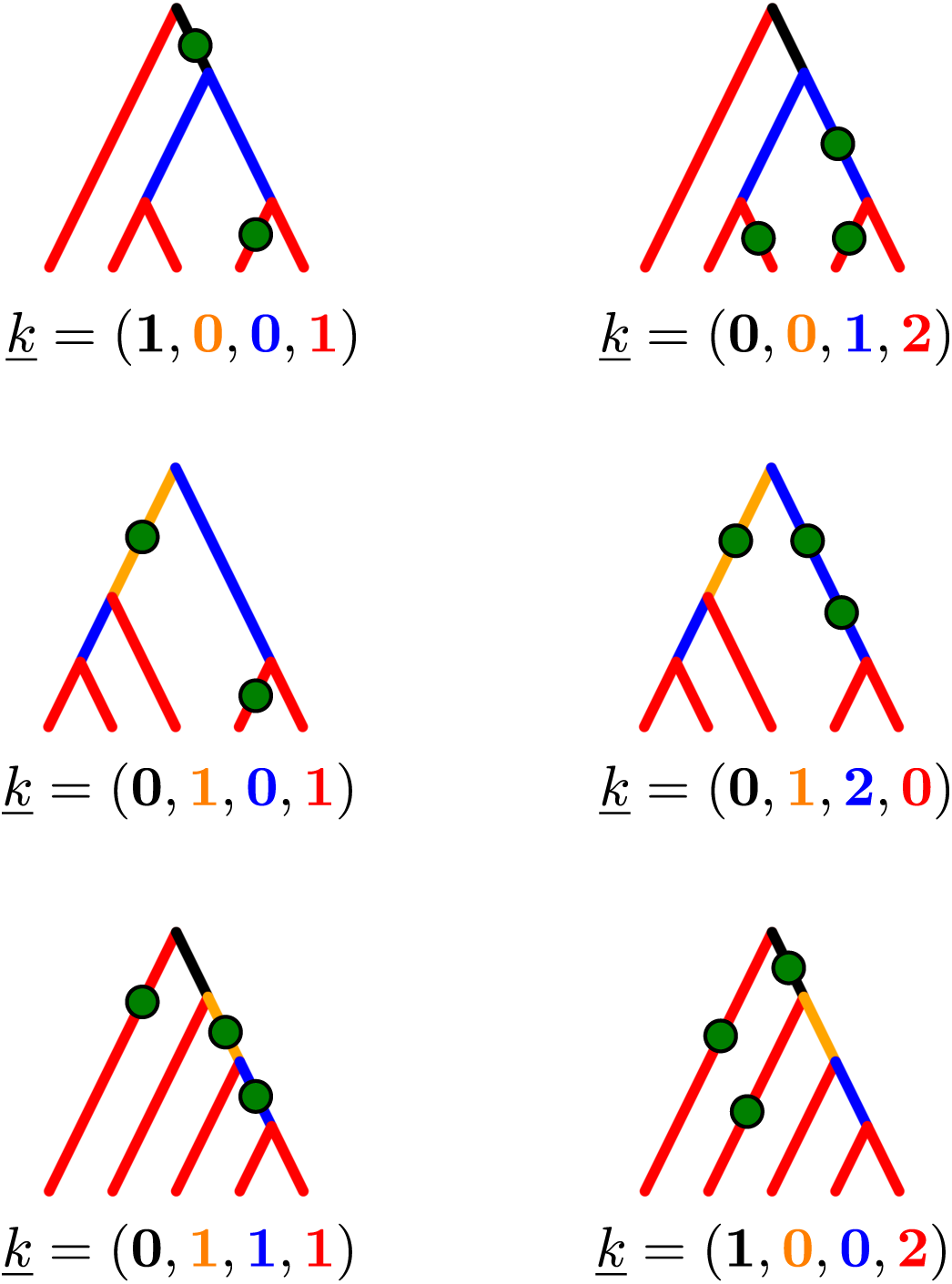
Example bSFS configurations for a sample from a single population. Genealogical relationships for a sample of size 5 can be generated by three types of topologies (top, middle and bottom rows). Ignoring information on the phase, the branches can be classified by the number of tips they are ancestral to; i.e. singletons (red), doubletons (blue), tripletons (orange) and quadrupletons (black). Further, mutations on these branches give rise to different bSFS configurations, i.e. vectors <u>*k*</u>.

**Fig. S2.**
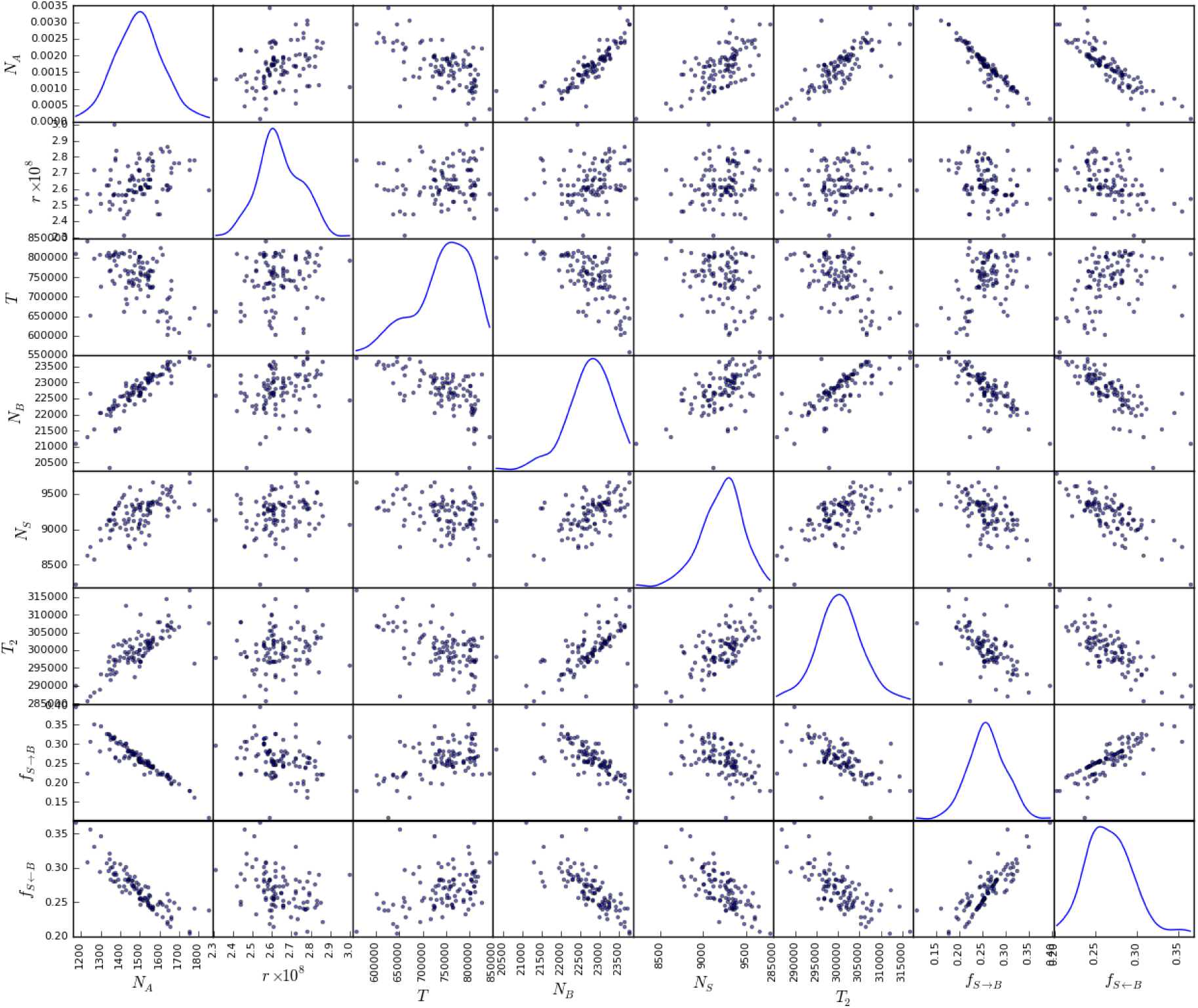
Correlations between the parametric bootstrap estimates obtained for the 2kb dataset (Table 2). The scatter plots in the lower and upper half of the figure (are symmetrically identical) show the pairwise correlations among parameter estimates under M6. Density plots for each parameter are respectively given along the diagonal.

**Fig. S3.**
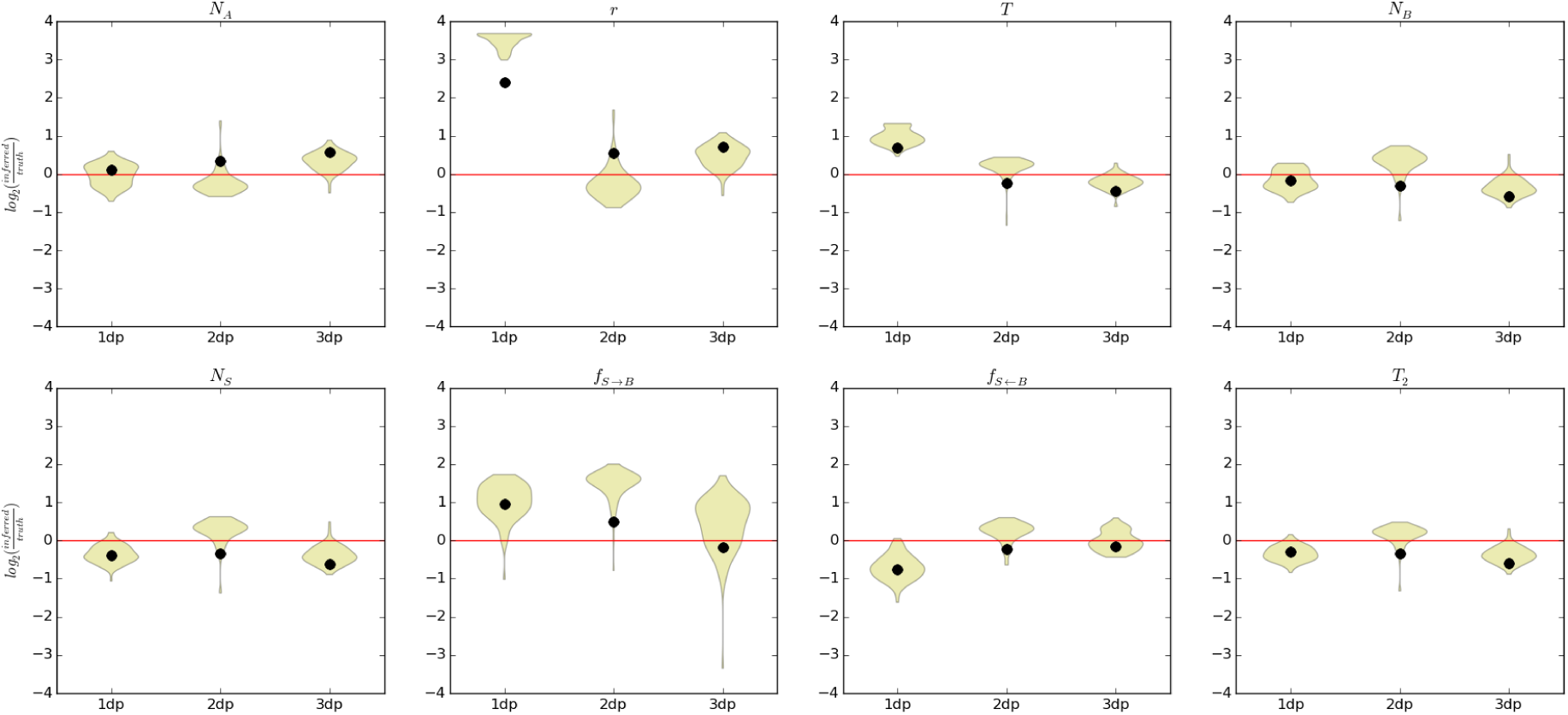
Parametric bootstrap results for model M6 and varying sample size. The *x*-axis indicates the number of diploid genomes per population. Accuracy for each parameter is given as the base-2 logarithm of the relative error (*i.e.* estimated value divided by the truth): 0 (red line) corresponds to the truth, increments of +1/-1 to two-fold over- or under-estimates. The black dots show the inferred point estimates (Table S3).

**Fig. S4.**
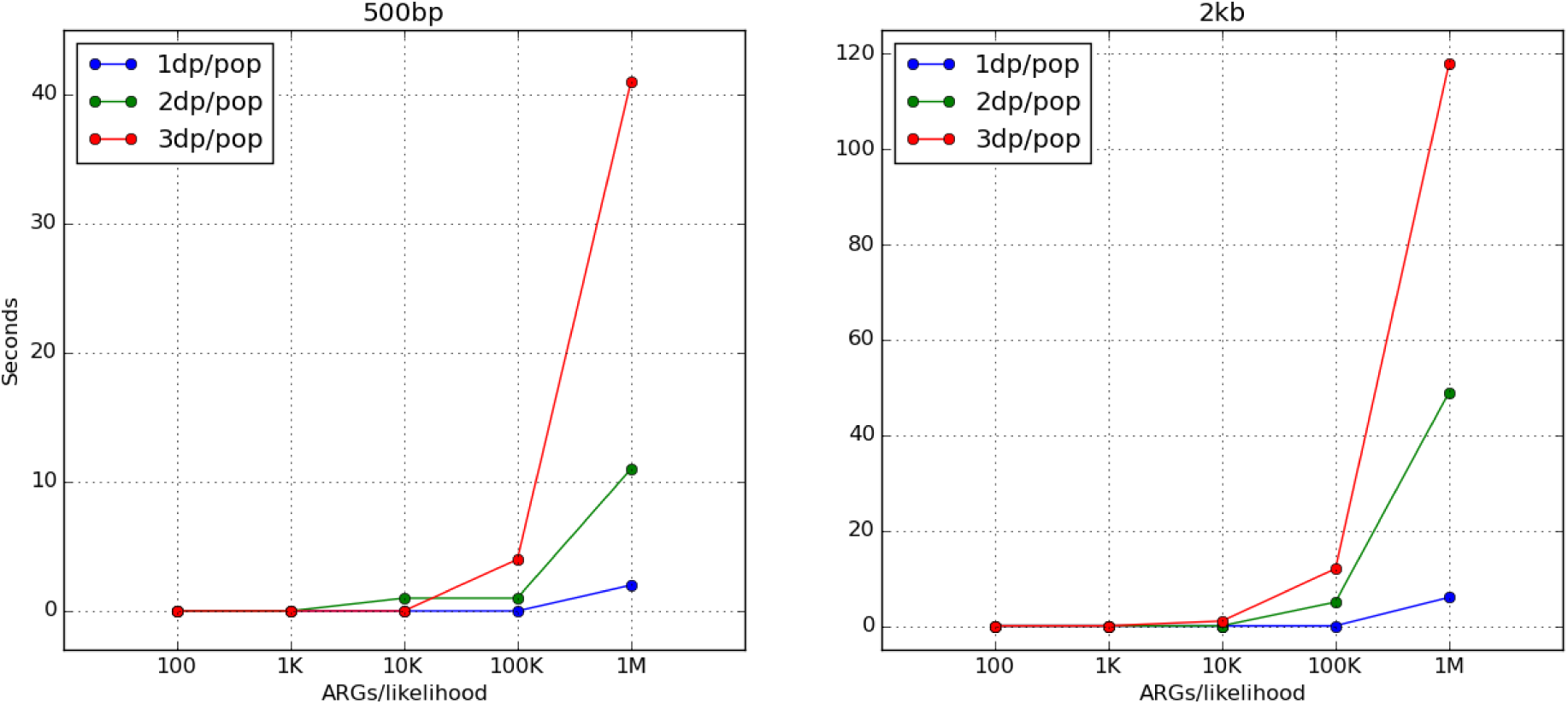
Computation times with varying sample and block sizes. Simulated datasets were generated by varying the number of diploid genomes per population under model M6 (for values used, see Table S3). Two types of block sizes were considered (500bp and 2kb) by keeping the whole genome size fixed at 200Mb. A single likelihood was evaluated at the true MCLE for each sampling configuration and block size by varying the number of sampled ARGs (*x*-axis). Times have been approximated to the nearest second.

**Fig. S5.**
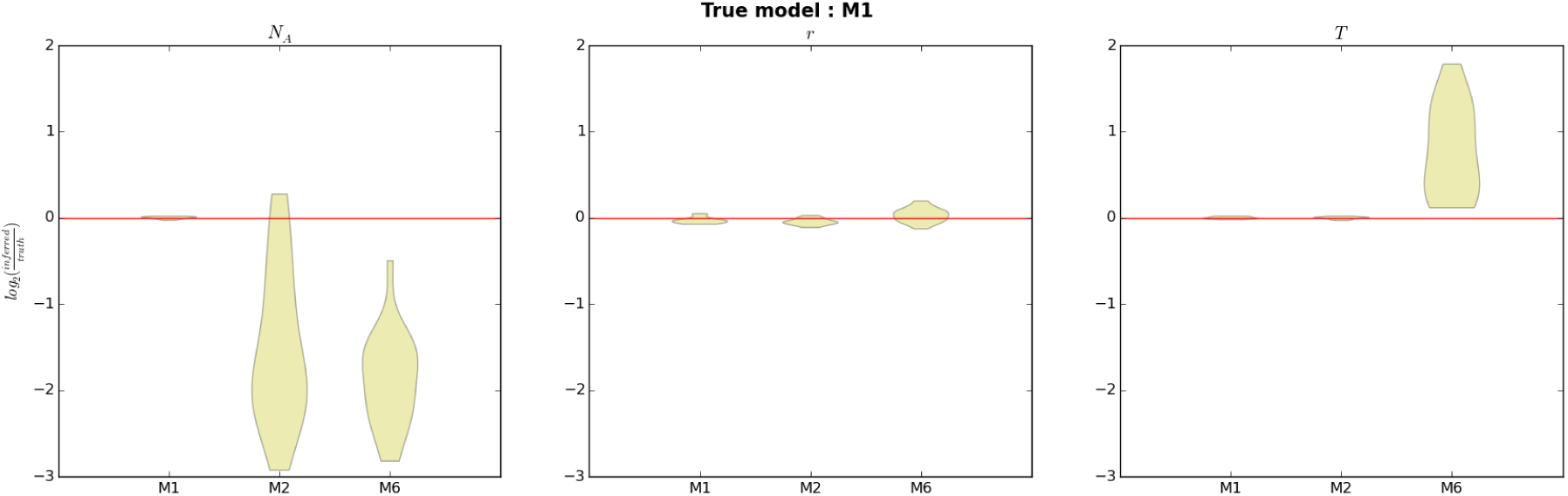
Parameter inference under the true and alternative models. The *y*-axis represents inference accuracy as given as the base-2 logarithm of the relative error (*i.e.* estimated value divided by the truth): 0 (red line) corresponds to the truth, increments of +1/-1 to two-fold over/under-estimates respectively.

**Fig. S6.**
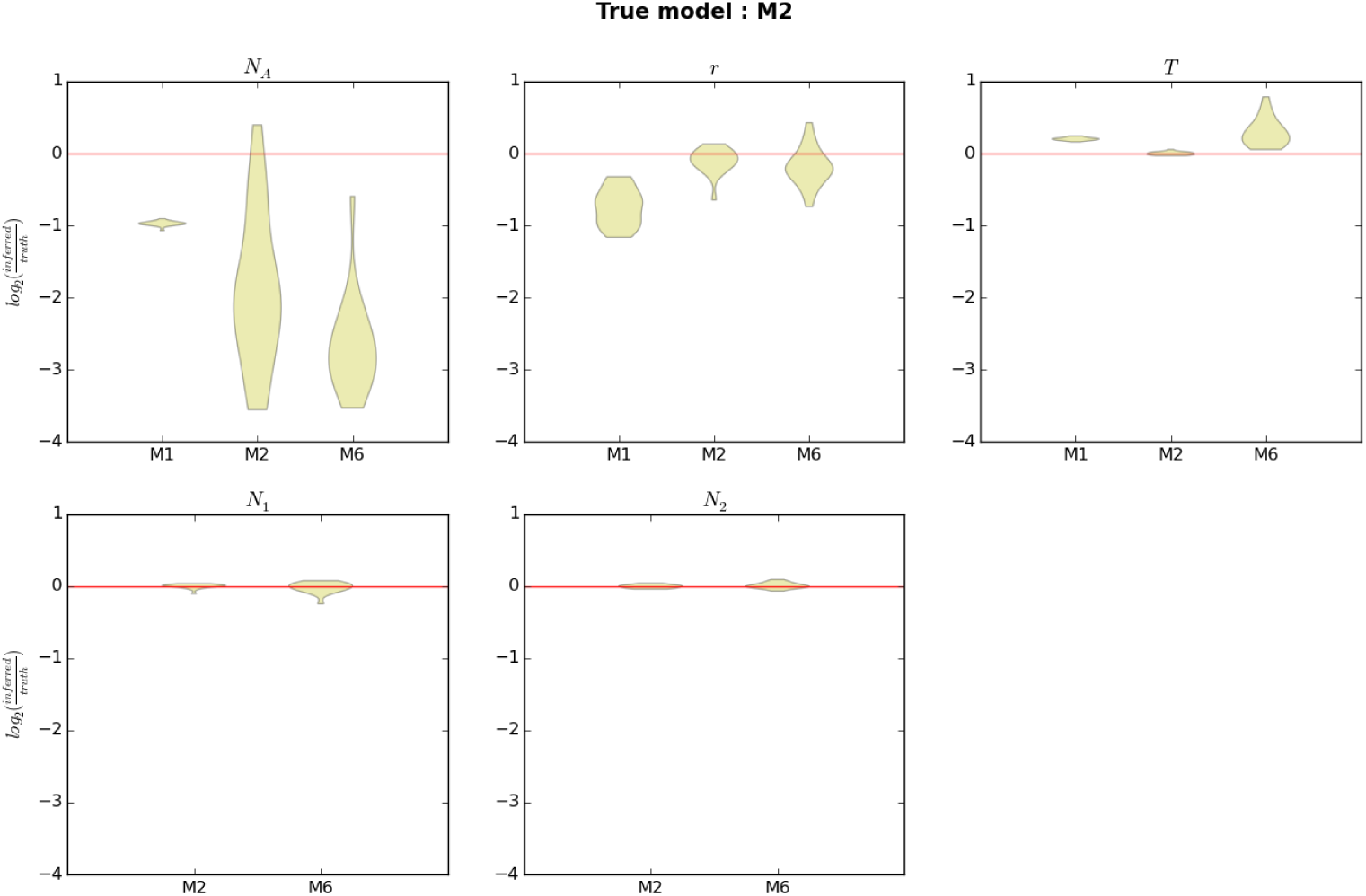
Parameter inference under the true and alternative models. The *y*-axis represents inference accuracy as given as the base-2 logarithm of the relative error (*i.e.* estimated value divided by the truth): 0 (red line) corresponds to the truth, increments of +1/-1 to two-fold over/under-estimates respectively.

**Fig. S7.**
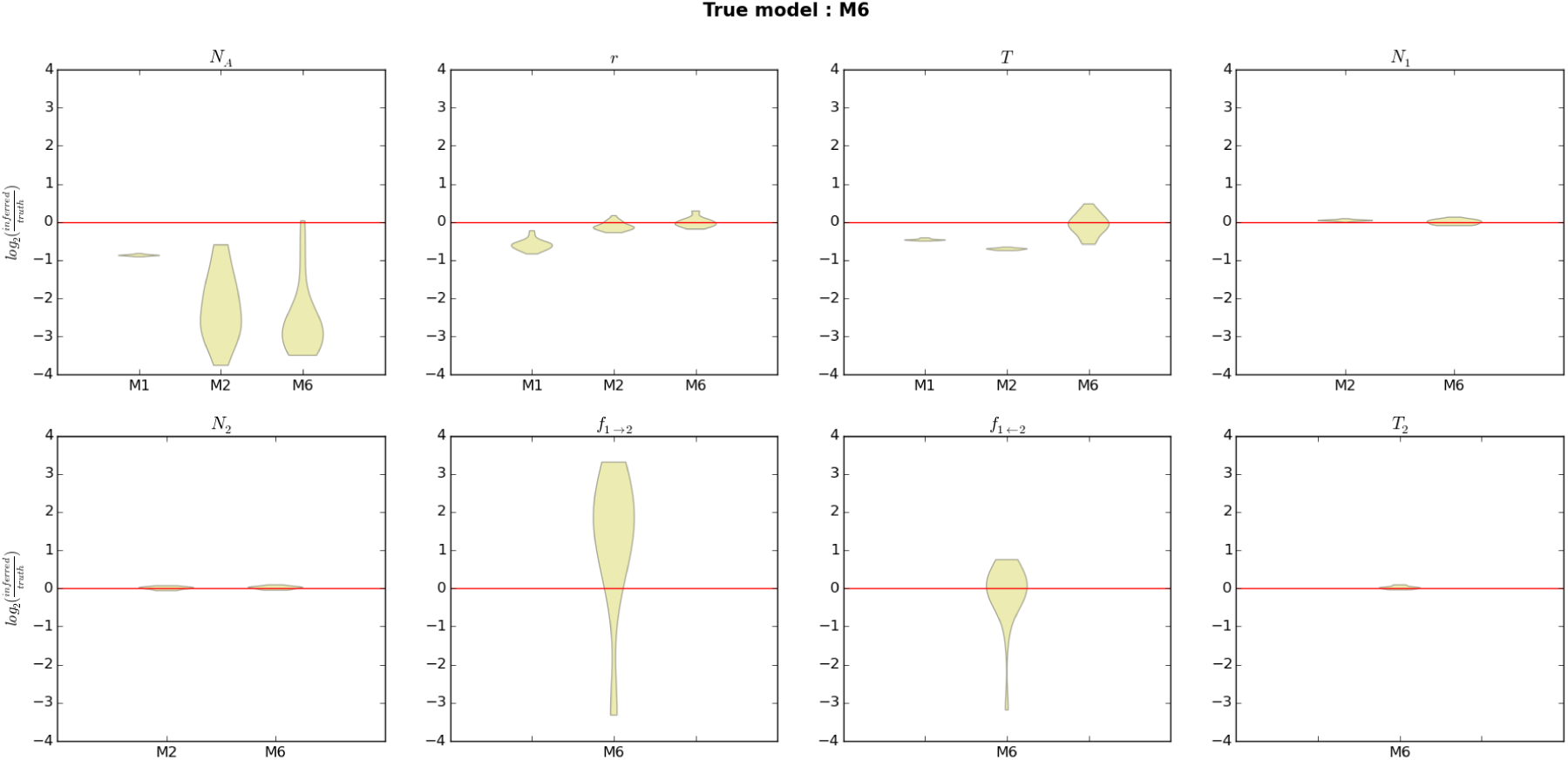
Parameter inference under the true and alternative models. The *y*-axis represents inference accuracy as given as the base-2 logarithm of the relative error (*i.e.* estimated value divided by the truth): 0 (red line) corresponds to the truth, increments of +1/-1 to two-fold over/under-estimates respectively.

**Fig. S8.**
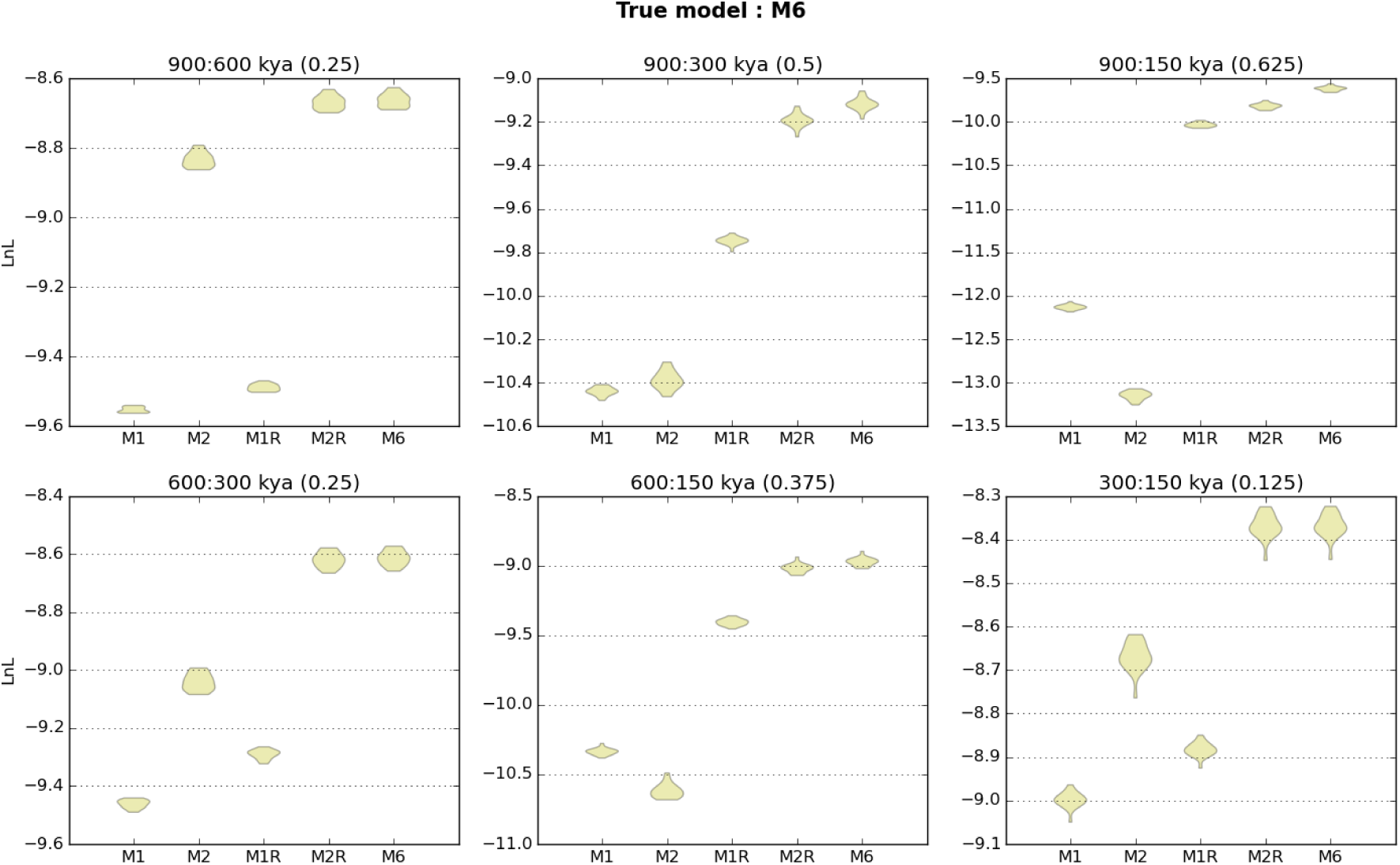
Nested model fit under the true (M6) and alternative models. Point LnLs were evaluated at the obtained MCLE under the true (M6) and alternative models and for the 20 simulated datasets/model. M1R & M2R represent alternative (to M1 & M2 respectively) scenarios where the divergence time between populations (*T*) was set to be that of the true admixture time(*i.e. T*_2_). Above each plot, the true split and admixture times are given in a *T*: *T*_2_ format and given in parenthesis is *T − T*_2_ in coalescent units. Higher LnL values (*i.e.* closer to zero) indicate a better fit to the simulated data.

**Fig. S9.**
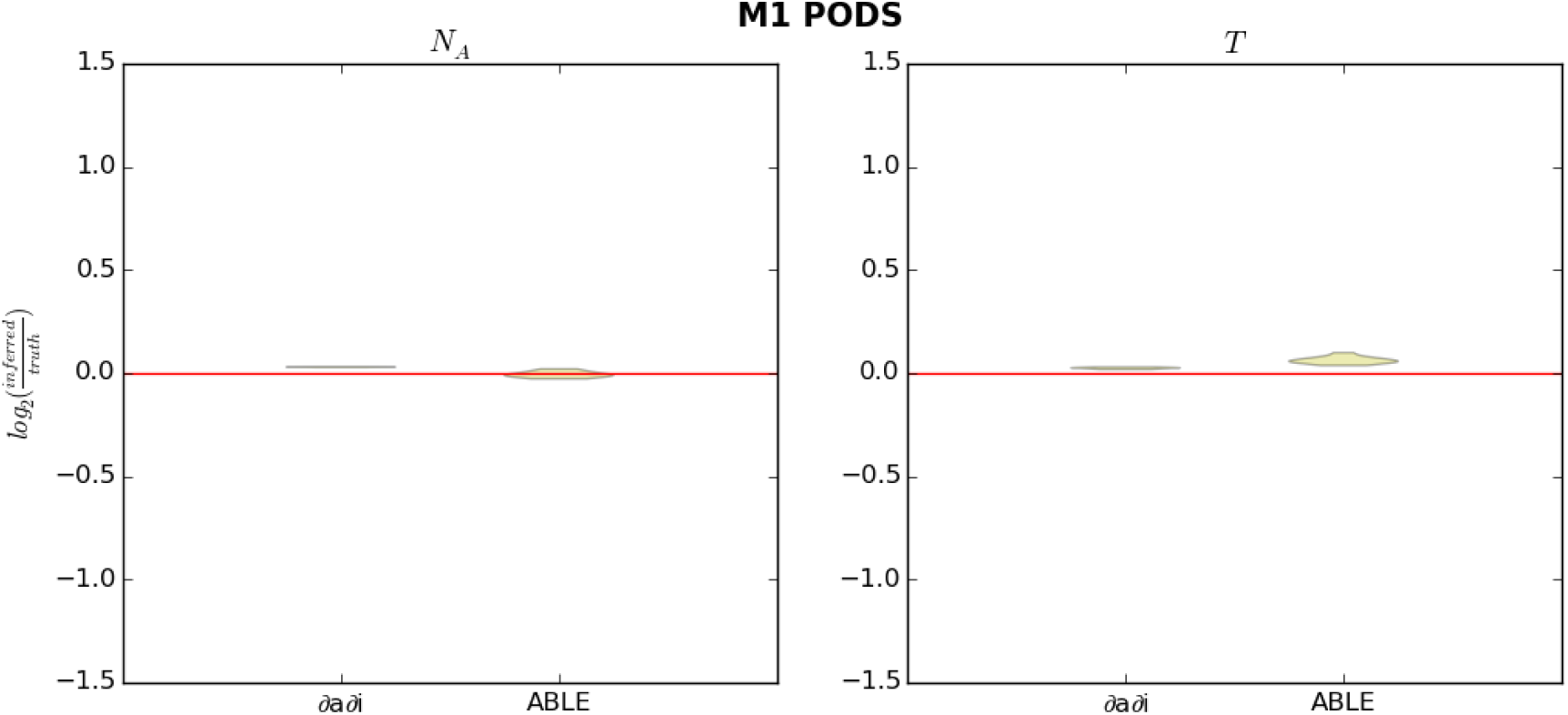
Comparison of ∂a∂i and ABLE under model M1. The *y*-axis represents inference accuracy as given as the base-2 logarithm of the relative error (*i.e.* estimated value divided by the truth): 0 (red line) corresponds to the truth, increments of +1/-1 to two-fold over/under-estimates respectively.

**Fig. S10.**
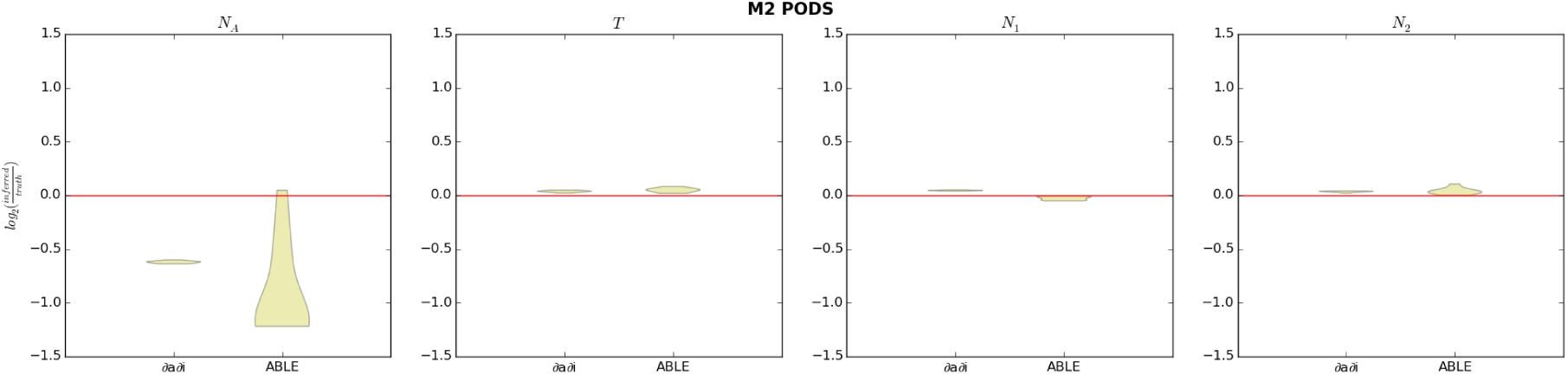
Comparison of ∂a∂i and ABLE under model M2. The *y*-axis represents inference accuracy as given as the base-2 logarithm of the relative error (*i.e.* estimated value divided by the truth): 0 (red line) corresponds to the truth, increments of +1/-1 to two-fold over/under-estimates respectively.

**Fig. S11.**
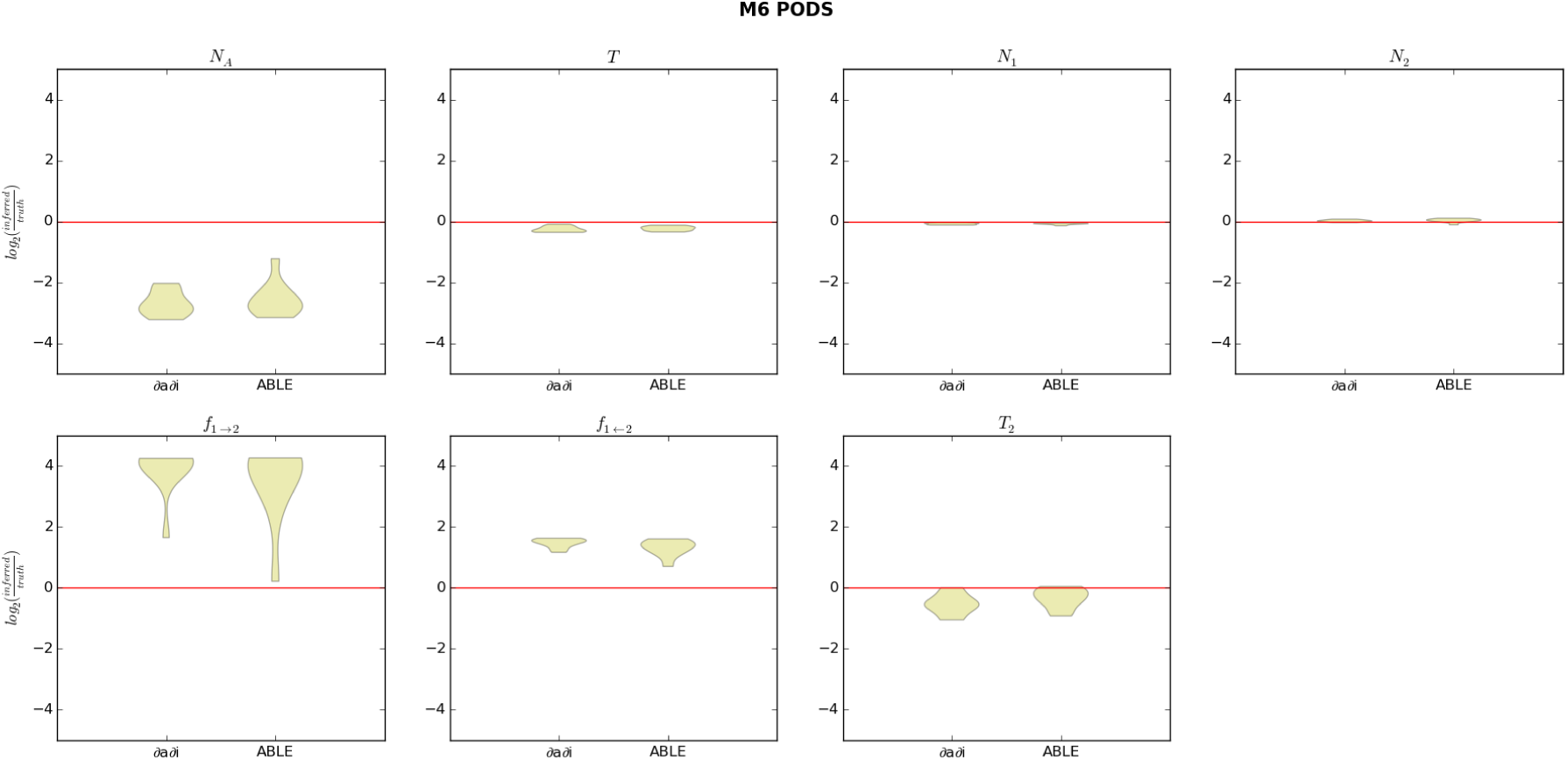
Comparison of ∂a∂i and ABLE under model M6. The *y*-axis represents inference accuracy as given as the base-2 logarithm of the relative error (*i.e.* estimated value divided by the truth): 0 (red line) corresponds to the truth, increments of +1/-1 to two-fold over/under-estimates respectively.

**Fig. S12.**
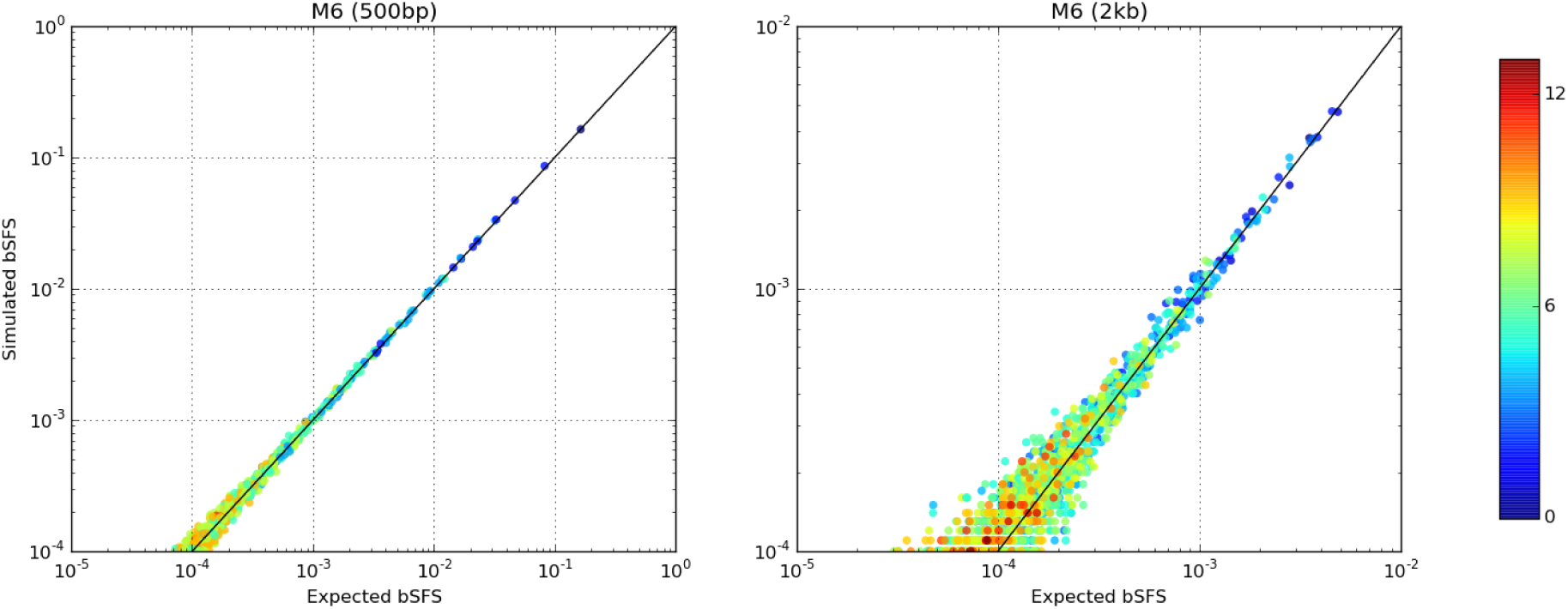
Effect of long range linkage on the absolute model fit using simulations and for the most common configurations. Two datasets were simulated under M6 using long (0.5 Mb) contiguous sequences totaling 200 Mb (cut into 500bp and 2kb blocks respectively) and with the inferred MCLEs from the orangutan dataset (Table S1 and Table 1 respectively). The expected bSFS (*x*-axis) was generated by sampling 50 million ARG’s at the inferred MCLEs and using the same values as for the simulated bSFS (*y*-axis). The diagonal black line indicates the perfect match between the expected and simulated. The colours represent the total number of SNPs contained in each configuration.

**Fig. S13.**
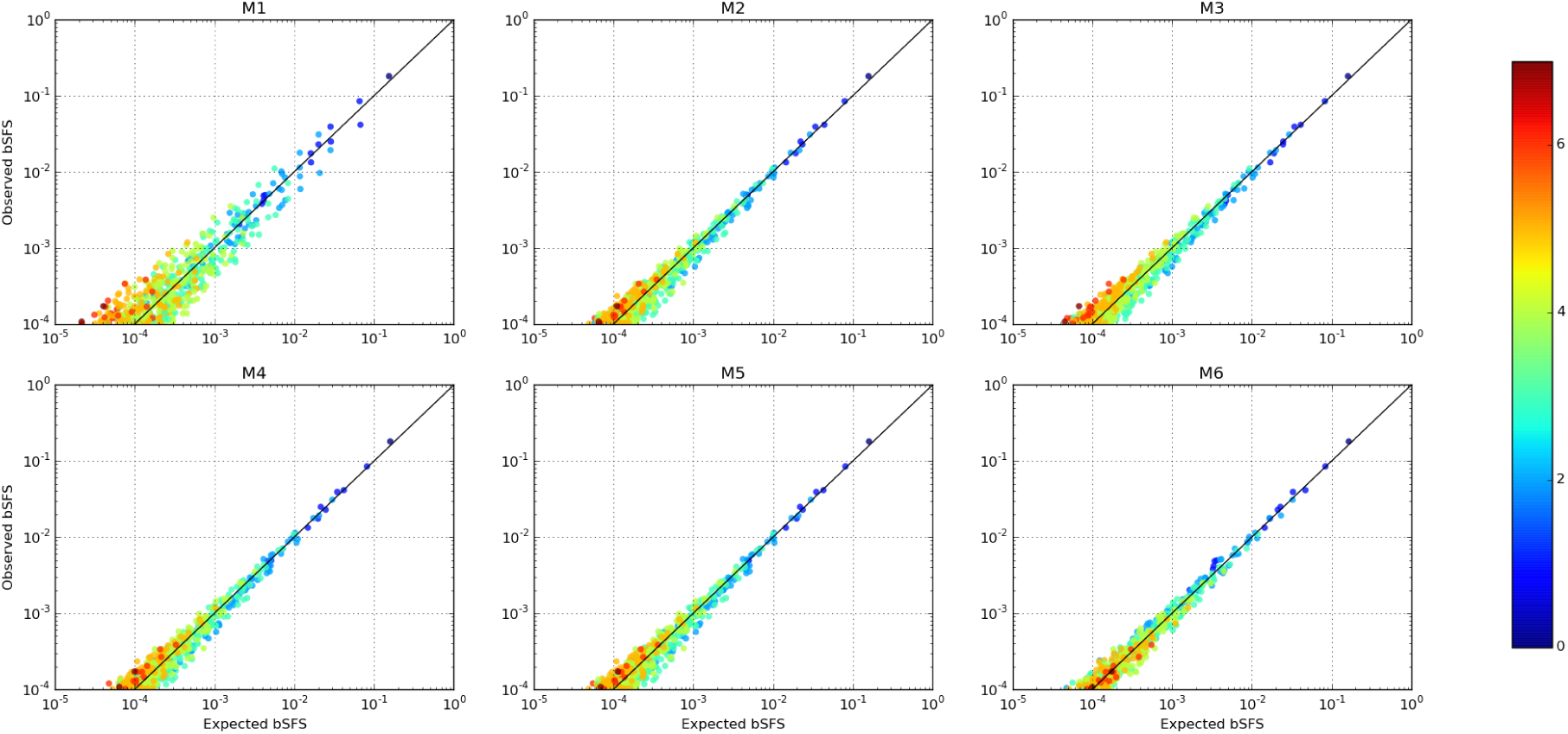
Absolute model fit to the observed 500bp bSFS for the most common configurations. Each point here represents a unique mutational configuration making up the bSFS. The expected bSFS (*x*-axis) was generated with ABLE using 50 million ARG’s at the MCLE for each model (Table S1) and plotted against the observed bSFS (*y*-axis) from the orangutan data. The diagonal black line indicates the perfect match between the expected and observed. The colours represent the total number of SNPs contained in each configuration.

**Fig. S14.**
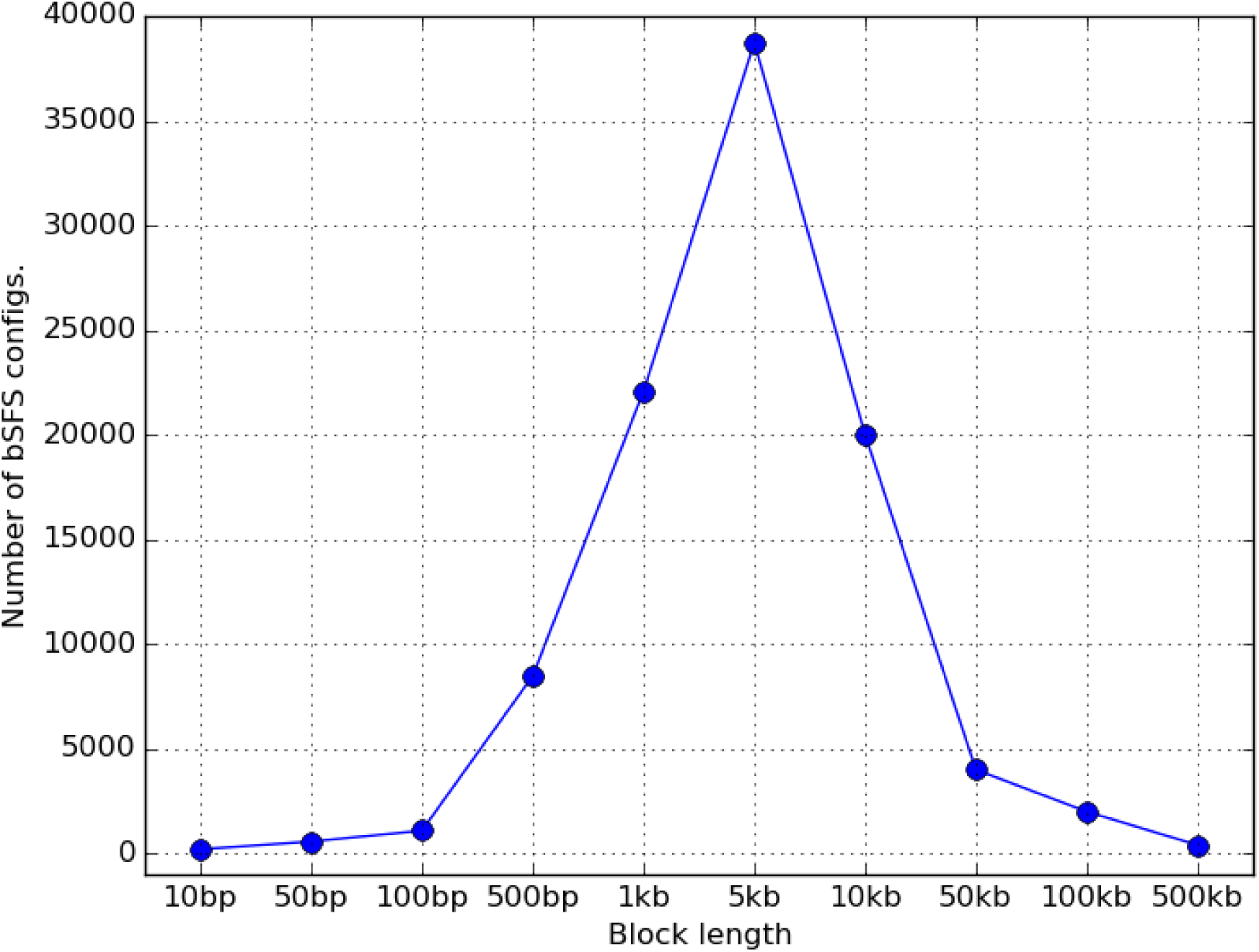
The number of unique bSFS configurations for different block sizes. Data sets of a total length of 200Mb sized were simulated as independent blocks of a fixed length (*x*-axis) under model M6 estimated from the 2kb MCLE data (see Table 1). For each block length the number of unique bSFS configurations is plotted on the *y*-axis. For this history, the number of configurations defined by the bSFS is maximized for blocks between 1kb - 10kb.

**Fig. S15.**
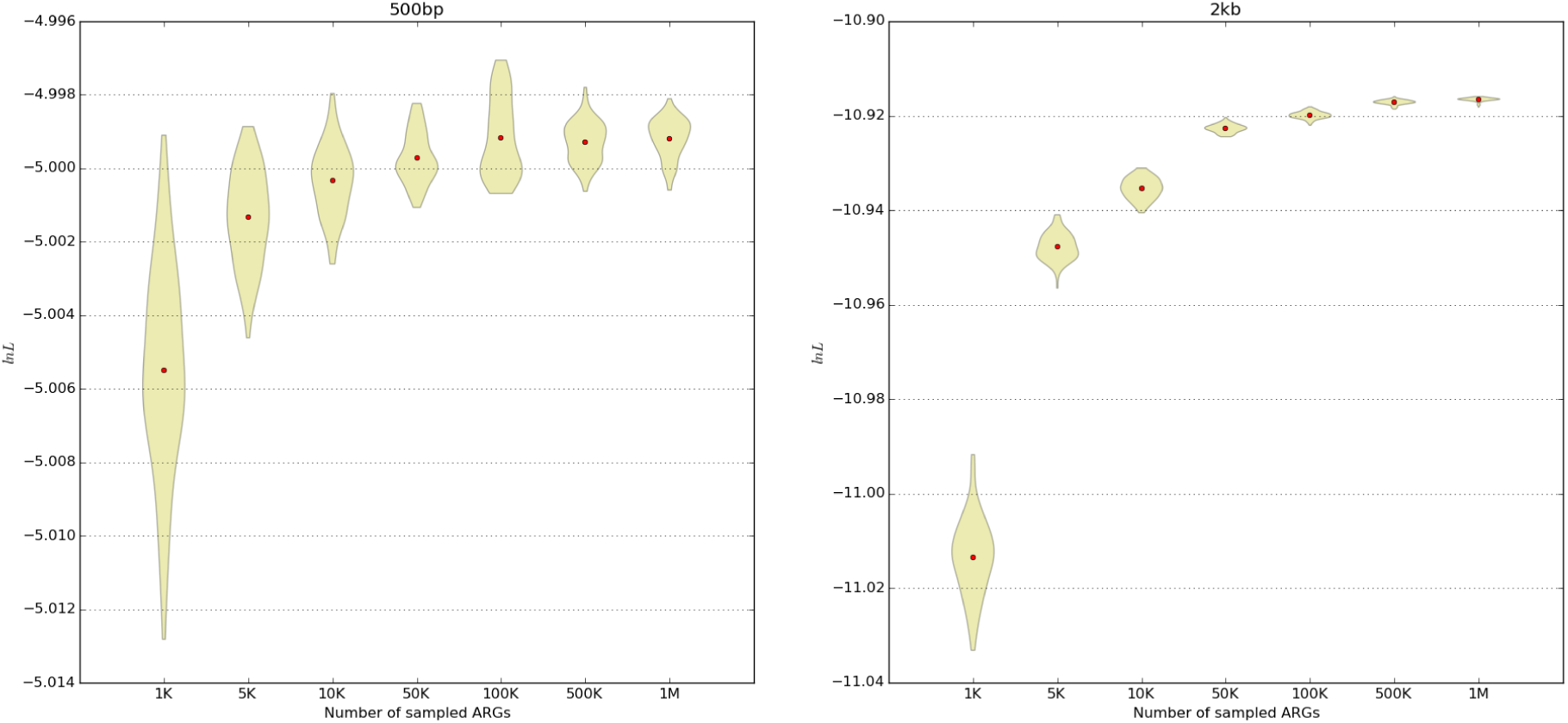
Accuracy of the log likelihood (LnL) approximation for M6. The LnL was evaluated at the MCLEs obtained for the 500bp (Table S1) and 2kb datasets (Table 1) for an increasing number of sampled ARGs (*x*-axis). Violin plots show the entire spread around the respective averages (red dots) over 100 log likelihood evaluations. Note that the *y*-axis represents the *per block* CLs *i.e.* downscaled with respect to the number of blocks found in each dataset.

**Fig. S16.**
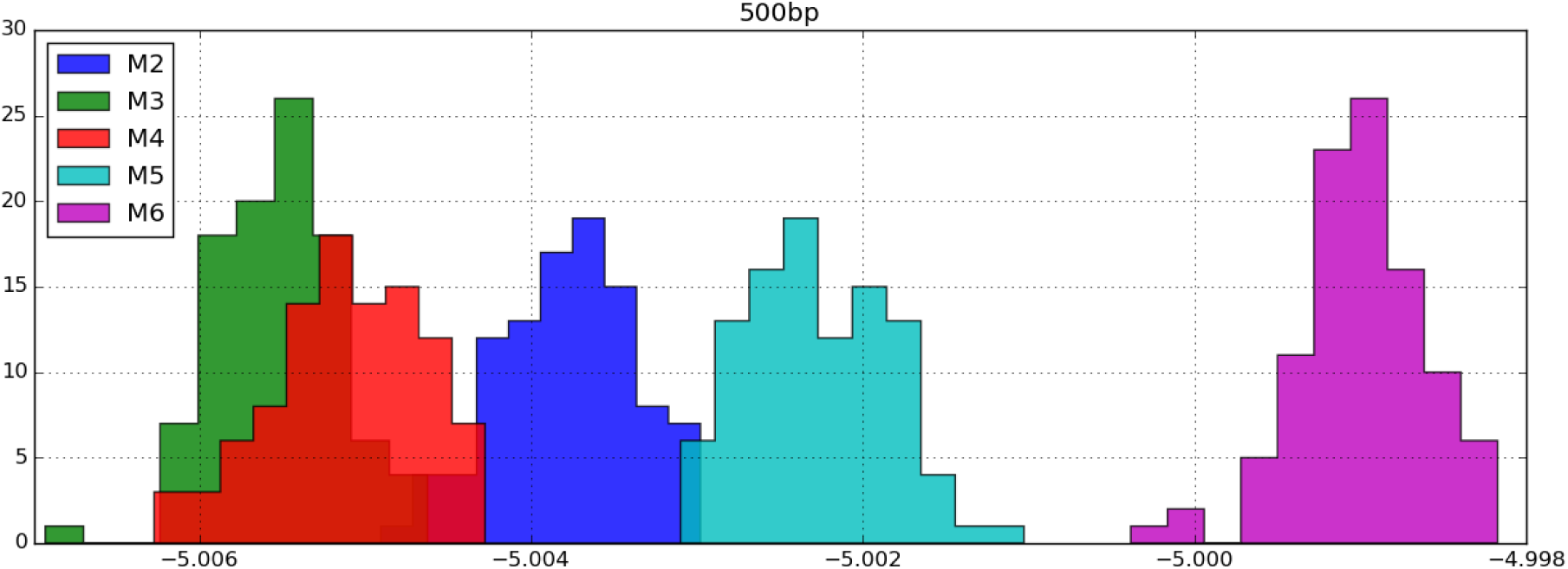
Relative fit between models using 500bp blocks. Histograms of 100 evaluations of the Composite Likelihood (CL) using 1 million ARG’s and at the MCLE corresponding to each model (Table Table S1). Note that the *x*-axis represents the *per block* CLs *i.e.* downscaled with respect to the number of blocks found in each dataset. Models further to the right are a relatively better fit than those to the left. Model M1 is ignored (appears much further to the left) as it has the worst fit among all models.

**Text. S1.**
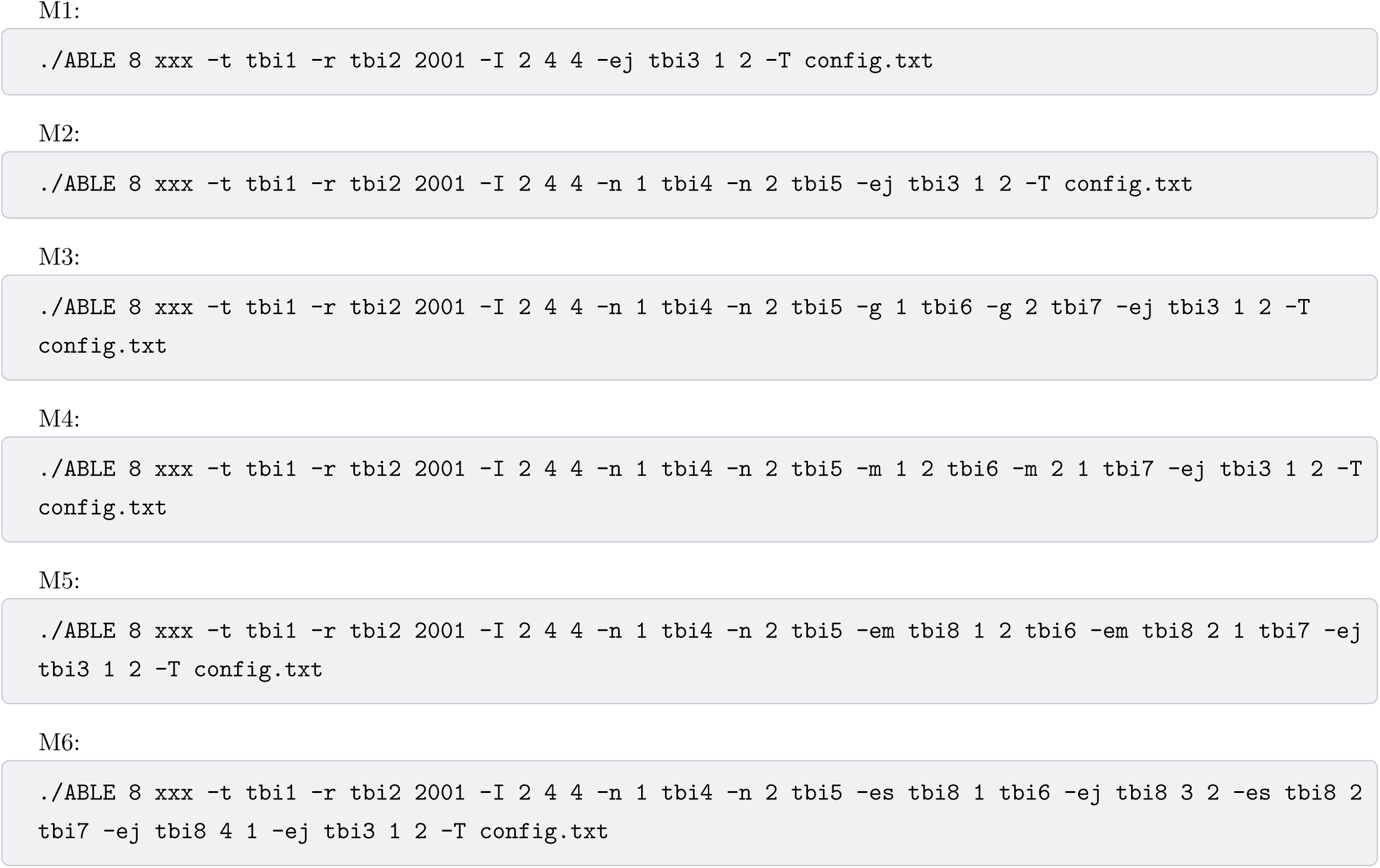
Command lines for launching models M1-M6 (see Fig. 4). Here we are assuming 2 diploid samples/population (which is also applicable to 4 haploid samples/population), block sizes of 2Kb and the working directory is that in which the ABLE binary is situated. Further documentation is available from https://github.com/champost/ABLE/raw/master/doc/helpABLE.pdf.

**Text. S2.**
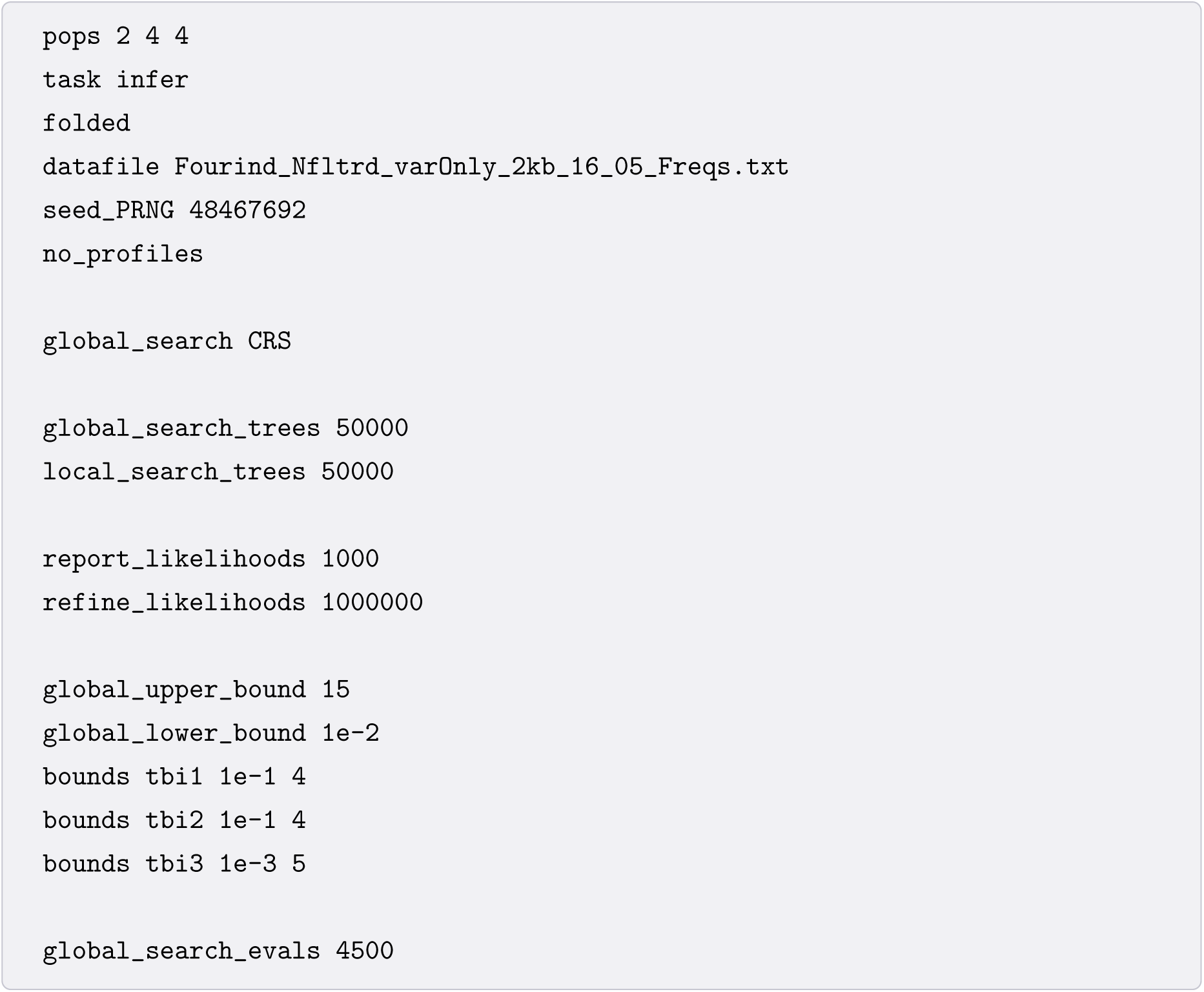
Configuration file for launching model M1 (see Fig. 4 and Text. S1). Here we are using 2Kb blocks from the Orangutan data consisting of 2 diploid samples each from the Bornean and Sumatran populations. The data files and further documentation is available for download from https://github.com/champost/ABLE/data.

**Text. S3.**
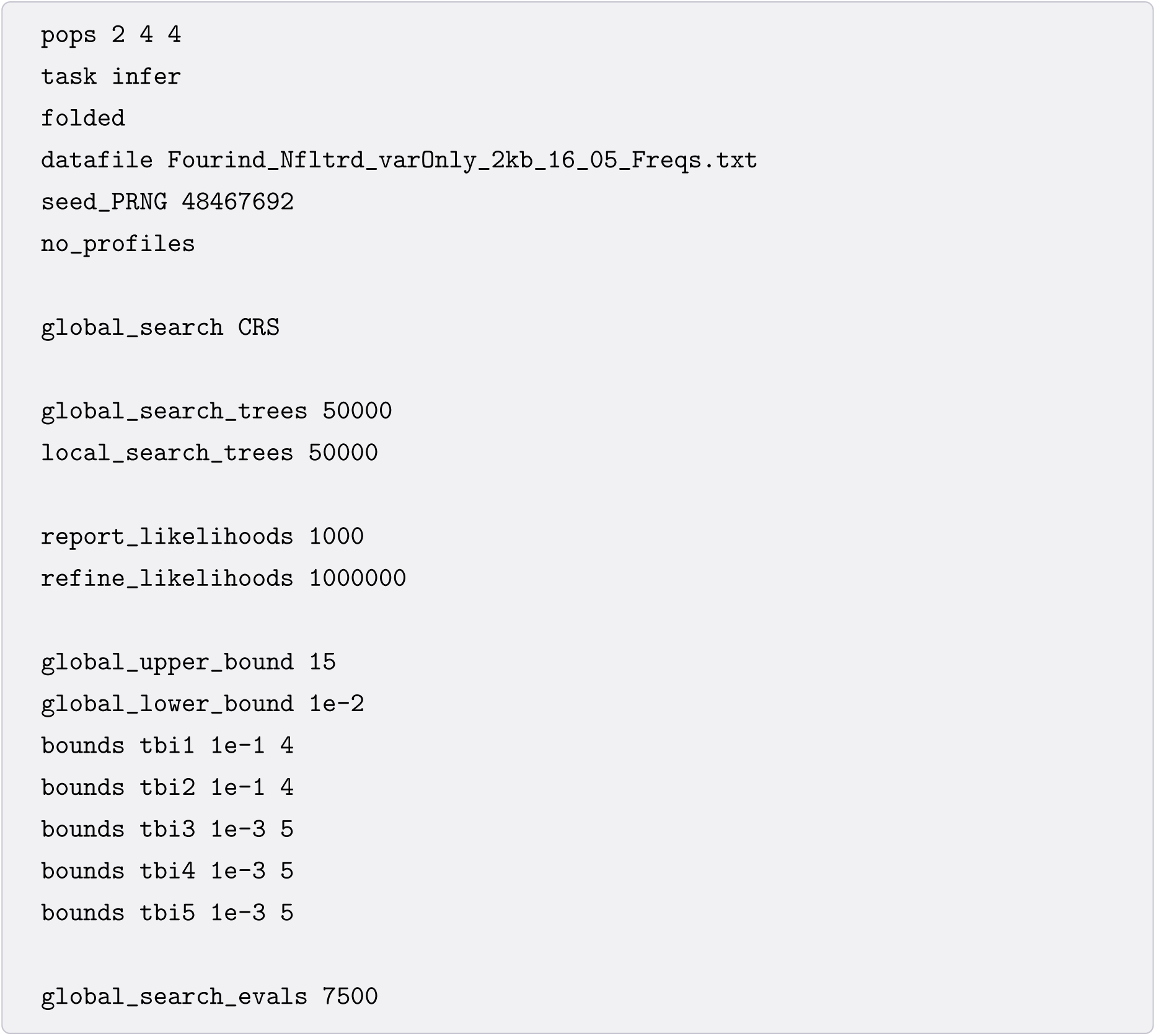
Configuration file for launching model M2 (see Fig. 4 and Text. S1). Here we are using 2Kb blocks from the Orangutan data consisting of 2 diploid samples each from the Bornean and Sumatran populations. The data files and further documentation is available for download from https://github.com/champost/ABLE/data.

**Text. S4.**
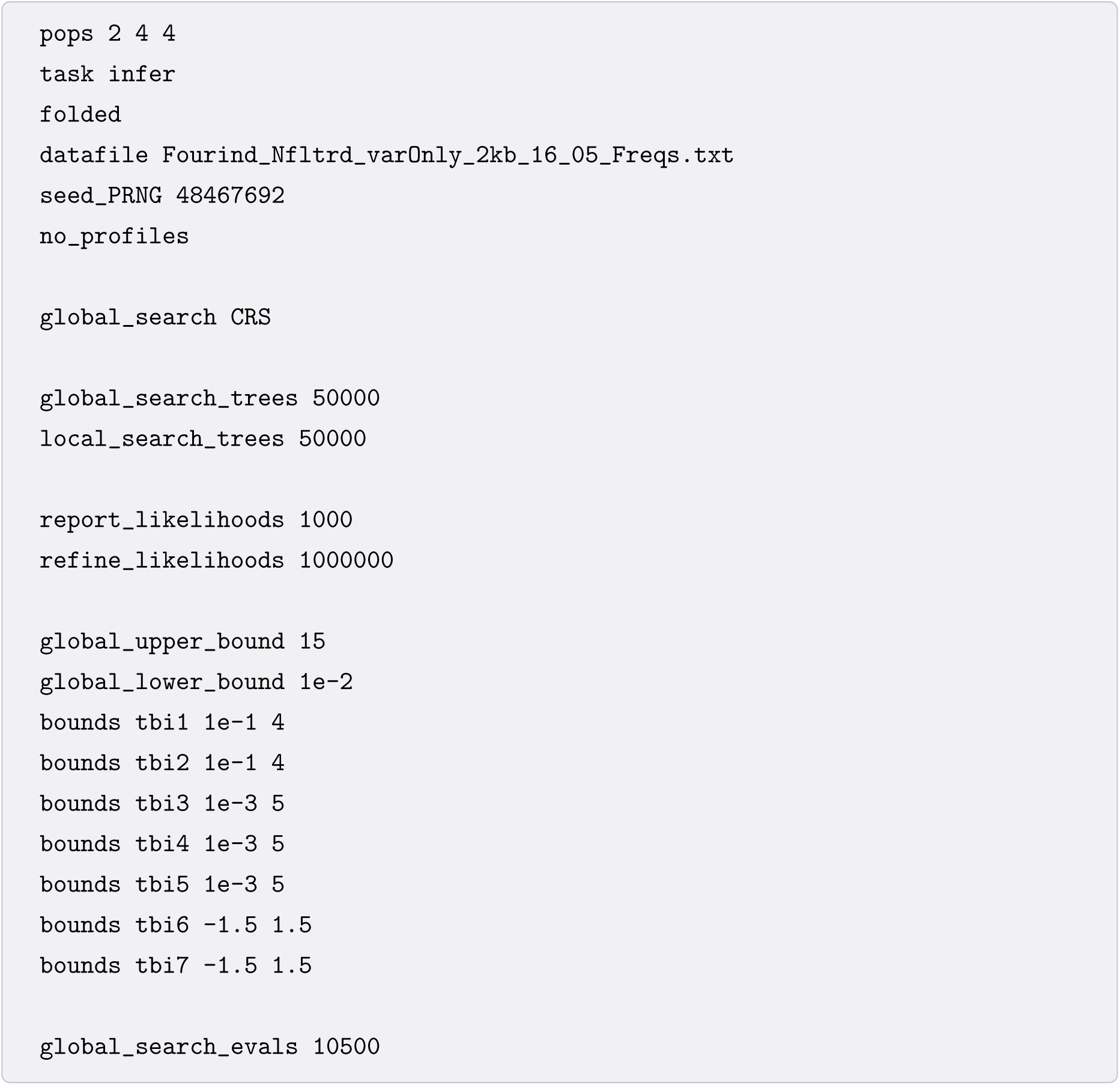
Configuration file for launching model M3 (see Fig. 4 and Text. S1). Here we are using 2Kb blocks from the Orangutan data consisting of 2 diploid samples each from the Bornean and Sumatran populations. The data files and further documentation is available for download from https://github.com/champost/ABLE/data.

**Text. S5.**
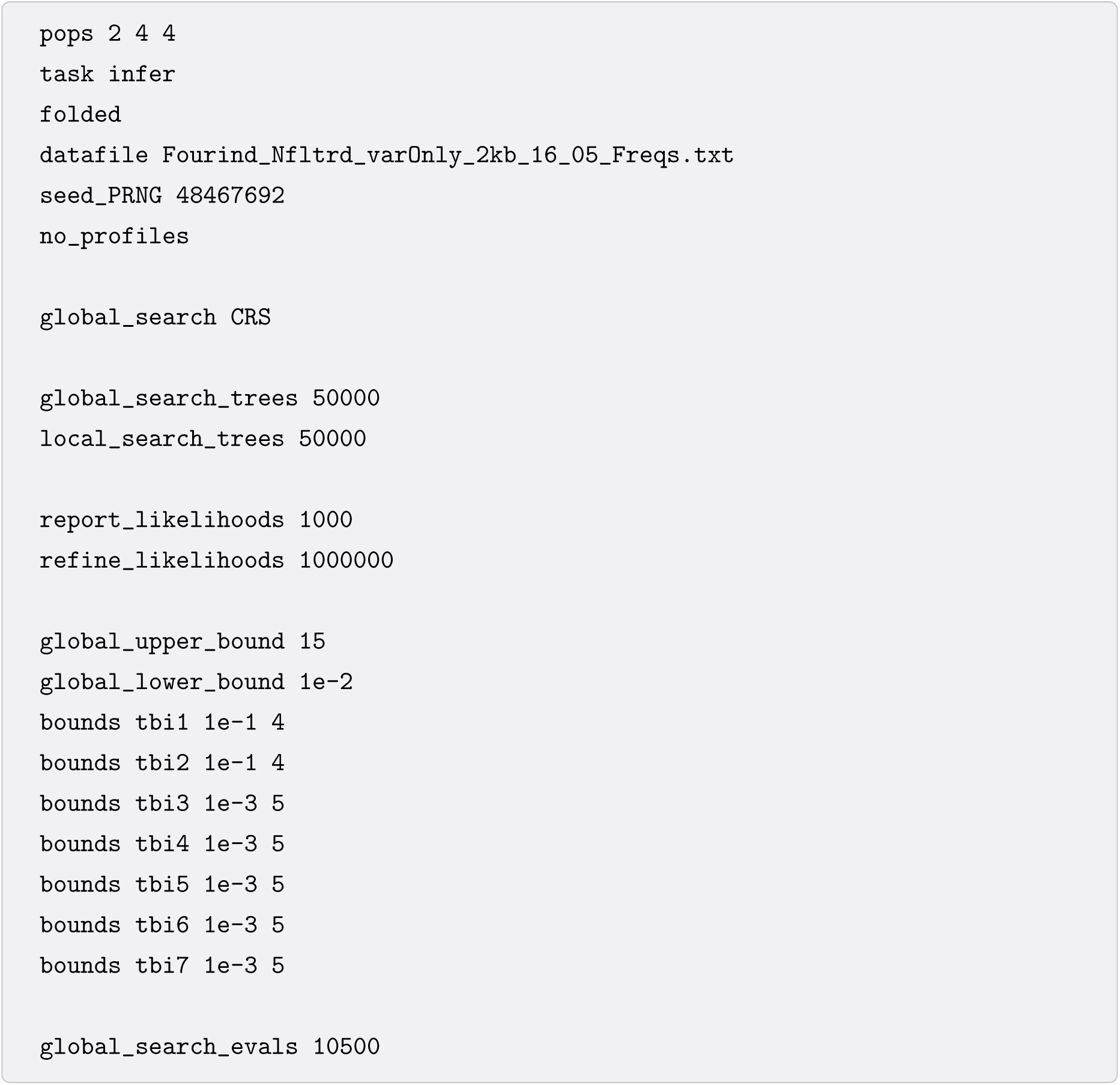
Configuration file for launching model M4 (see Fig. 4 and Text. S1). Here we are using 2Kb blocks from the Orangutan data consisting of 2 diploid samples each from the Bornean and Sumatran populations. The data files and further documentation is available for download from https://github.com/champost/ABLE/data.

**Text. S6.**
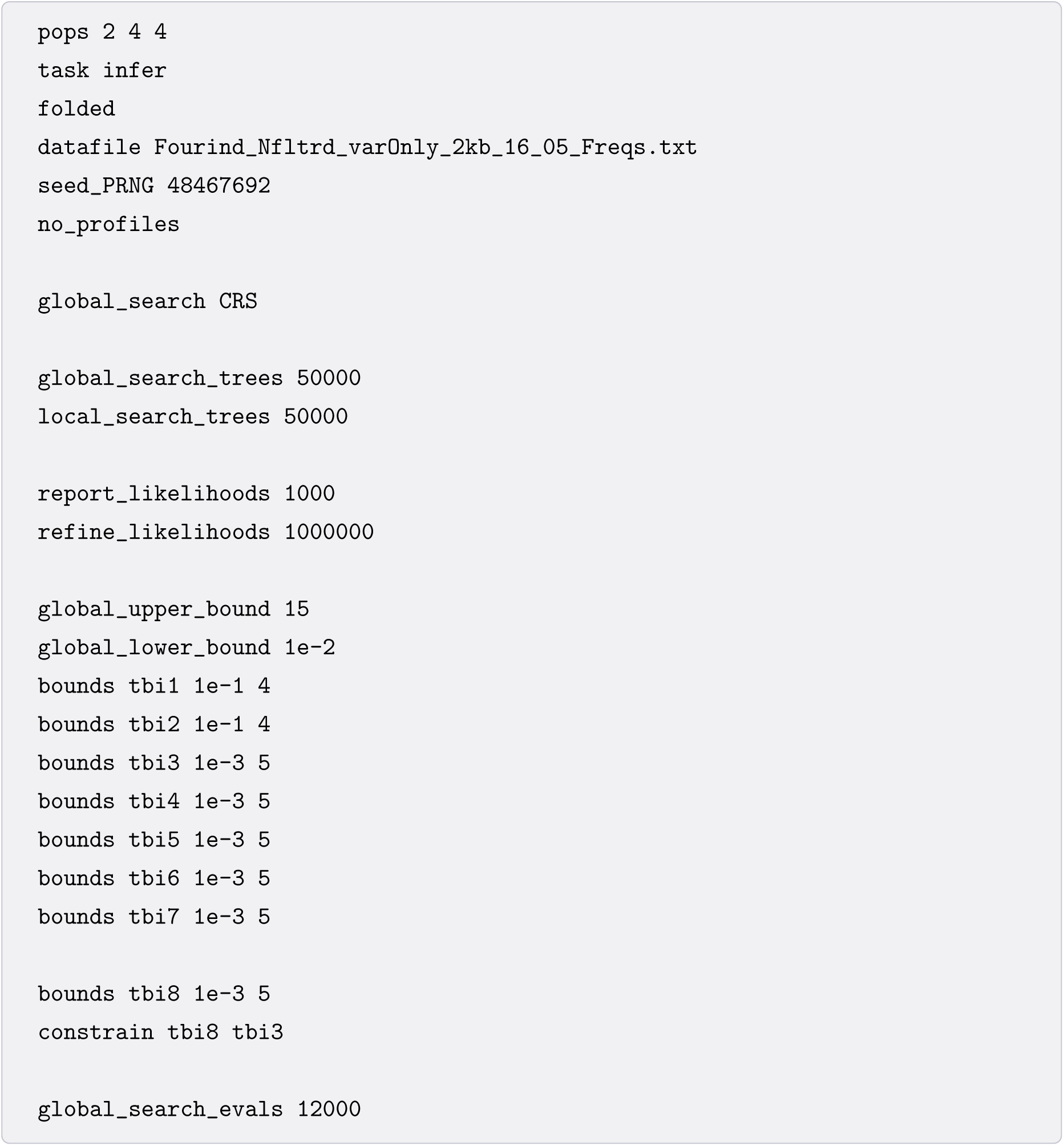
Configuration file for launching model M5 (see Fig. 4 and Text. S1). Here we are using 2Kb blocks from the Orangutan data consisting of 2 diploid samples each from the Bornean and Sumatran populations. The data files and further documentation is available for download from https://github.com/champost/ABLE/data.

**Text. S7.**
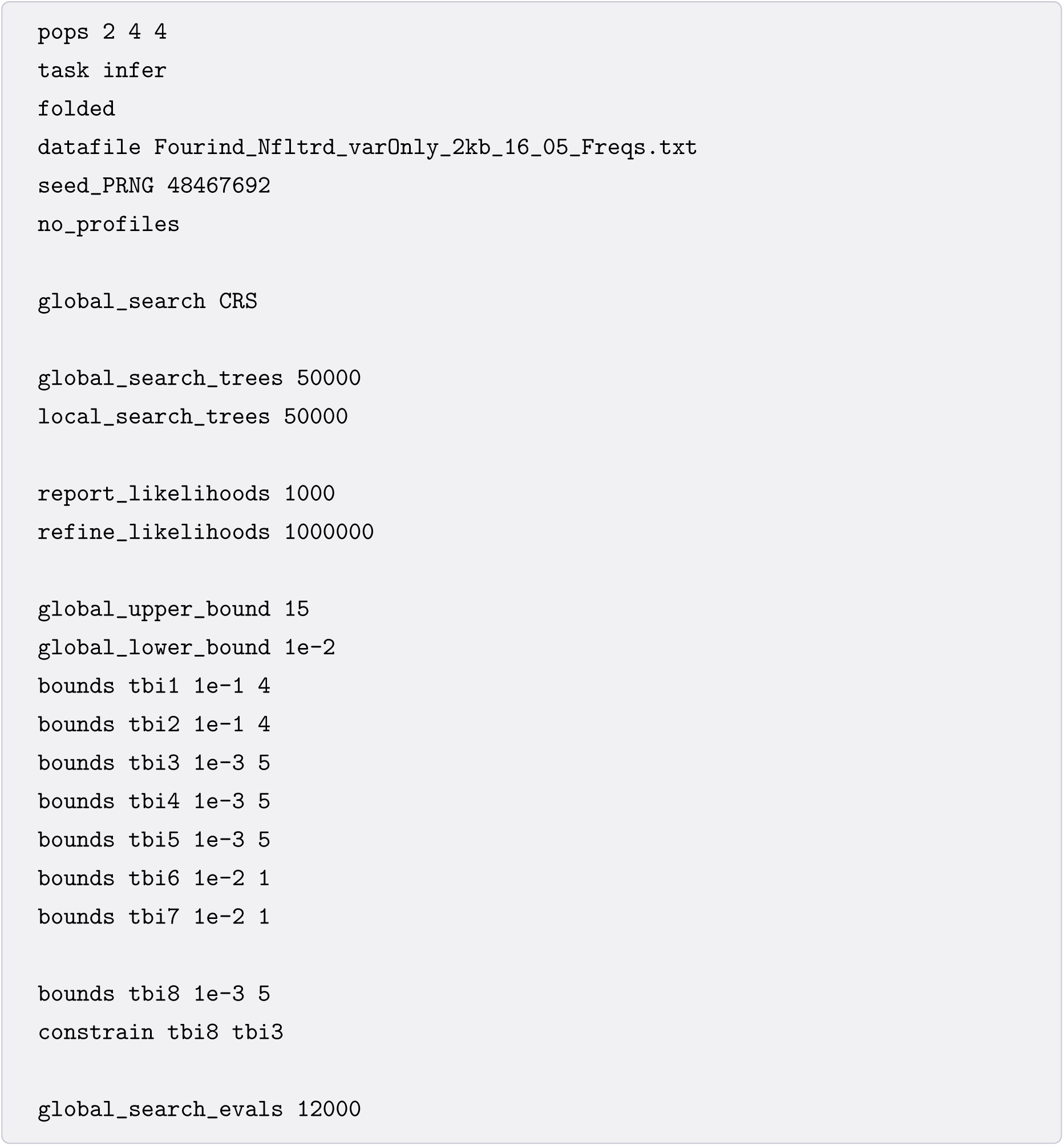
Configuration file for launching model M6 (see Fig. 4 and Text. S1). Here we are using 2Kb blocks from the Orangutan data consisting of 2 diploid samples each from the Bornean and Sumatran populations. The data files and further documentation is available for download from https://github.com/champost/ABLE/data.

**Text. S8.**
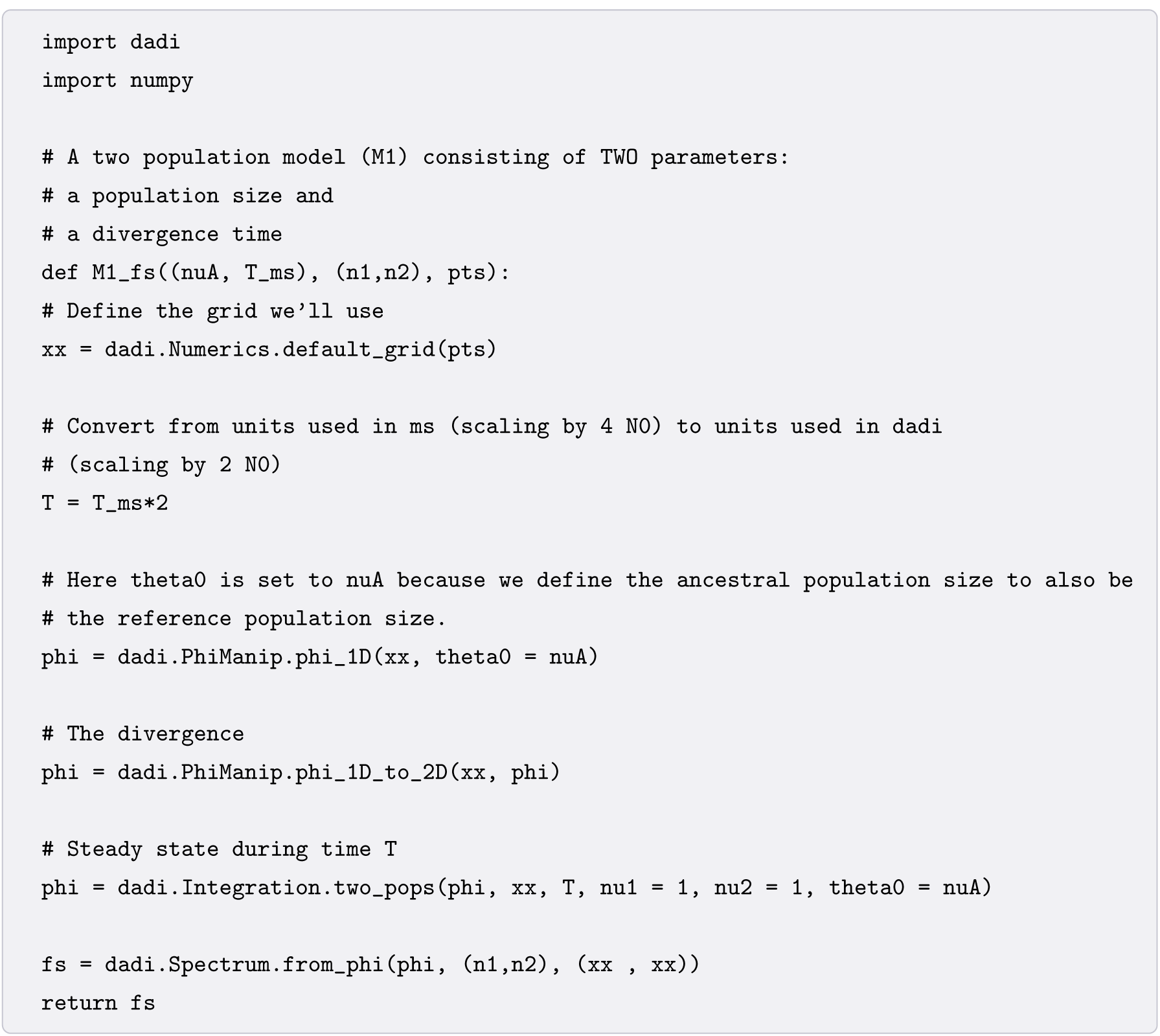
Python code (for a ∂a∂i analysis) defining model M1 (see Fig. 4). The divergence time parameters have been expressed in *ms* [41] units for convenience. This code can be imported into your main python script which calls the ∂a∂i inference procedure.

**Text. S9.**
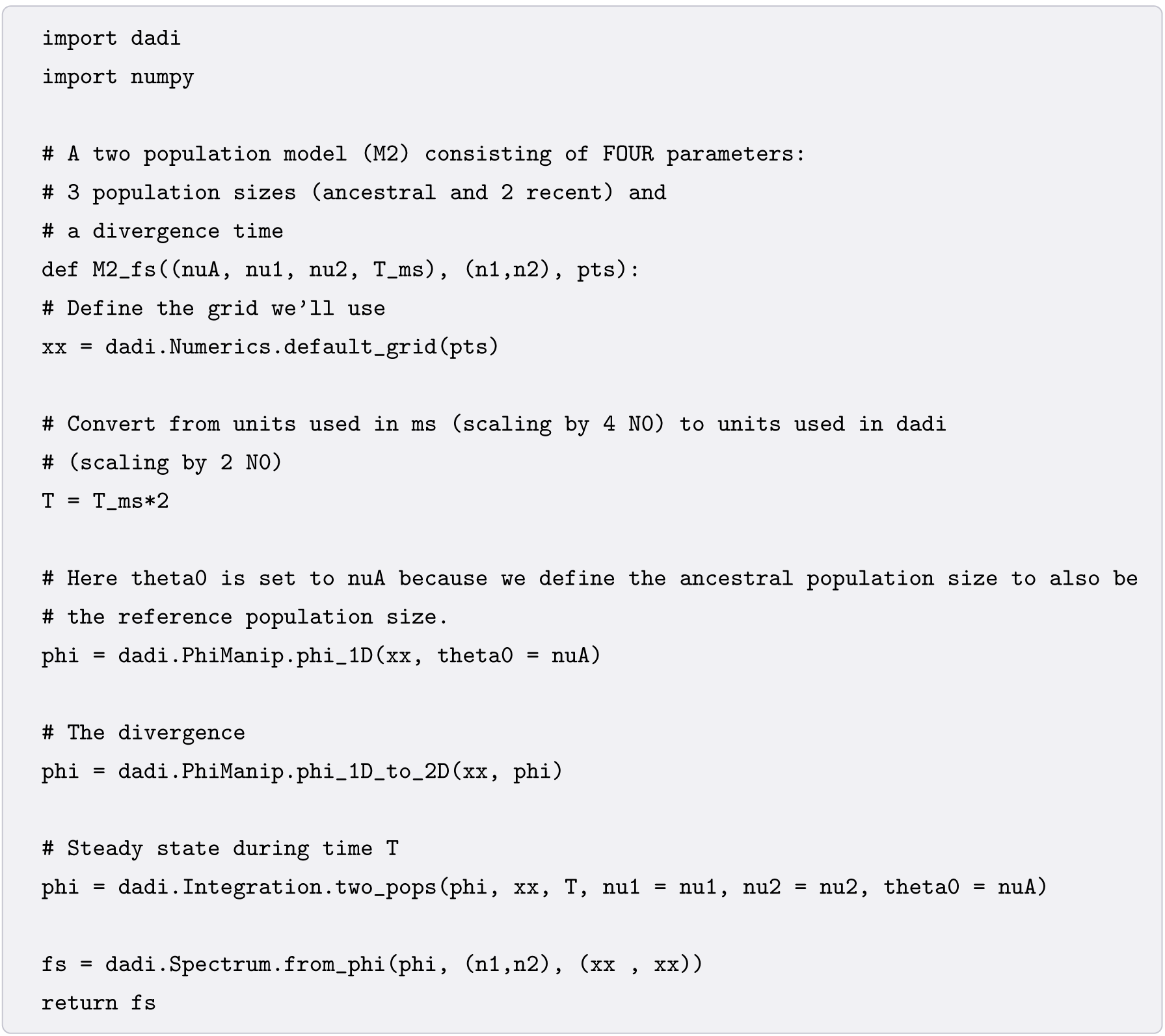
Python code (for a ∂a∂i analysis) defining model M2 (see Fig. 4). The divergence time parameters have been expressed in *ms* [41] units for convenience. This code can be imported into your main python script which calls the ∂a∂i inference procedure.

**Text. S10.**
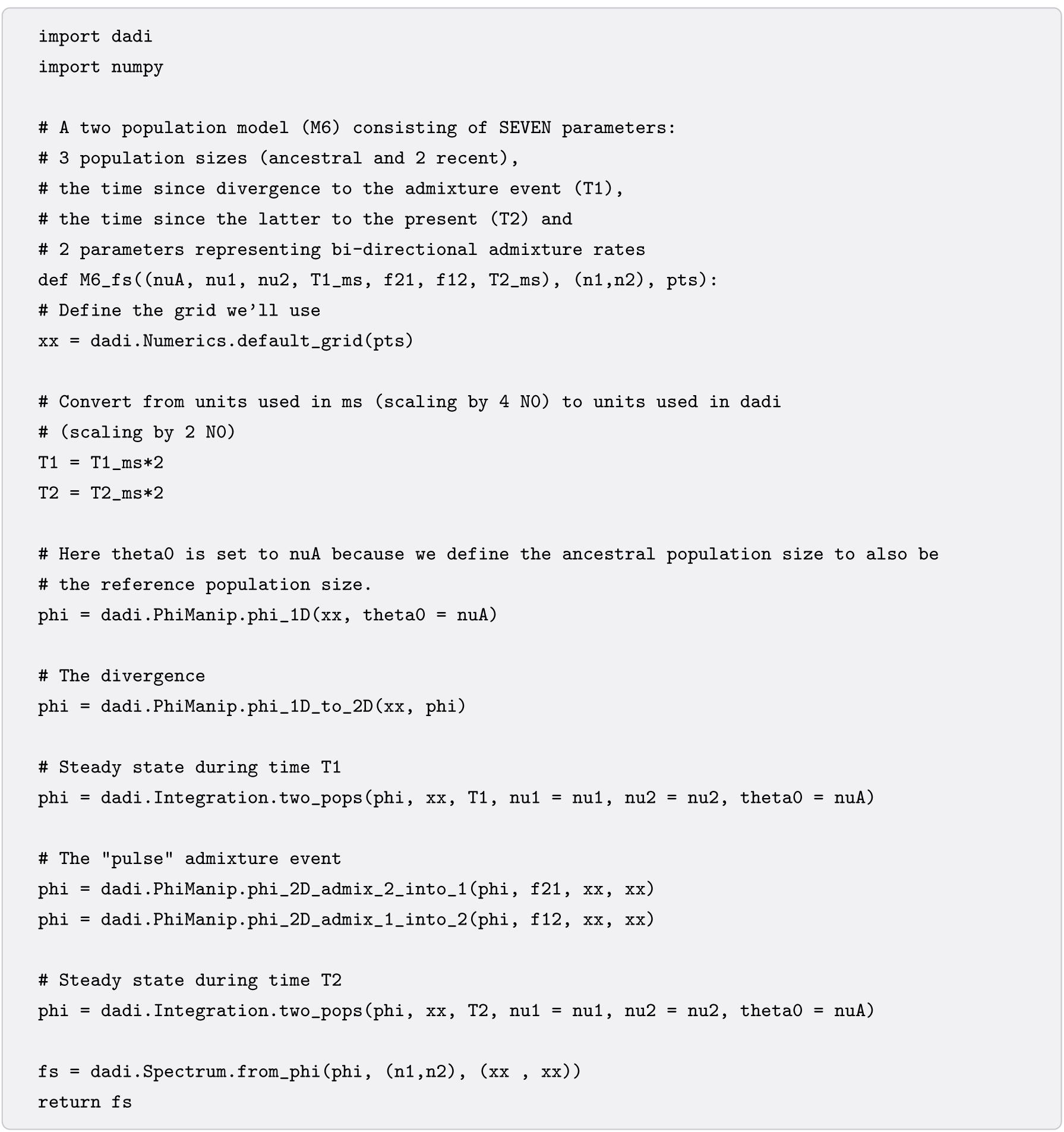
Python code (for a ∂a∂i analysis) defining models M6 (see Fig. 4). The divergence and admixture time parameters have been expressed in *ms* [41] units for convenience. This code can be imported into your main python script which calls the ∂a∂i inference procedure.

